# Histone acetylation in an Alzheimer’s disease cell model promotes homeostatic amyloid-reducing pathways

**DOI:** 10.1101/2023.09.18.558276

**Authors:** Daniel C. Xu, Hanna Sas-Nowosielska, Greg Donahue, Hua Huang, Naemeh Pourshafie, Charly R. Good, Shelley L. Berger

## Abstract

Alzheimer’s Disease (AD) is a disorder characterized by cognitive decline, neurodegeneration, and accumulation of amyloid plaques and tau neurofibrillary tangles in the brain. Dysregulation of epigenetic histone modifications may lead to expression of transcriptional programs that play a role either in protecting against disease genesis or in worsening of disease pathology. One such histone modification, acetylation of histone H3 lysine residue 27 (H3K27ac), is primarily localized to genomic enhancer regions and promotes active gene transcription. We previously discovered H3K27ac to be more abundant in AD patient brain tissue compared to the brains of age-matched non-demented controls. In this study, we use iPSC-neurons derived from familial AD patients with an amyloid precursor protein (APP) duplication (APP^Dup^ neurons) as a model to study the functional effect of lowering CBP/P300 enzymes that catalyze H3K27ac primarily at gene enhancers. We found that homeostatic amyloid-reducing genes were upregulated in the APP^Dup^ neurons compared to non-demented controls. We lowered CBP/P300 to reduce H3K27ac, which led to decreased expression of numerous of these homeostatic amyloid-reducing genes, along with increased extracellular secretion of a toxic amyloid-β species, Aβ(1-42). Our findings suggest that epigenomic histone acetylation, including H3K27ac, drives expression of compensatory genetic programs in response to AD-associated insults, specifically those resulting from APP duplication, and thus may play a role in mitigating AD pathology in neurons.

## Introduction

Alzheimer’s Disease (AD), the most common form of senile dementia, is a neurodegenerative disease characterized clinically by irreversible memory loss and cognitive decline^1^. The pathology of AD is comprised mainly of amyloid-β (Aβ) plaques^2–5^ and tau neurofibrillary tangles^6^. However, the relationship of these pathological hallmarks to each other and to disease features such as cognitive decline and neurodegeneration remain incompletely understood^7,8^.

Advances in understanding of chromatin architecture dynamics during normal aging and during neurodegeneration have revealed epigenomic dysregulation as a potential important link connecting aging, disease pathological features, and neurodegeneration^9^. In one example, we and others have uncovered a disease-association for histone 3 acetylation at lysine residue 27 (H3K27ac), a histone modification catalyzed by the lysine acetyltransferases p300 (EP300) and CBP (CREBBP)^10^. H3K27ac is found primarily in enhancer regions, and promotes active transcription of genes^11–13^. However, the balance of epigenomic H3K27ac gains and losses (genomic peaks measured using ChIPseq) is less clear, and indeed may be brain region-dependent and gene-dependent. Specifically, while we observed a substantial net increase in the number of H3K27ac peaks in the lateral temporal lobe of AD patients^14^, there was a net decrease in H3K27ac peaks in the entorhinal cortex (although a substantial number of H3K27ac peaks were still gained in AD patient samples compared to control)^15^. Hence, the precise occurrence and role of enhancer-associated H3K27ac may be complex and gene-specific, and requires additional clarification.

Efforts to determine a functional relationship, whether disease-promoting or disease-ameliorating, between altered H3K27ac abundance and pathology are complicated by the complexity of brains and the difficulty in establishing causality in postmortem tissue samples, where pathology and epigenomic dysregulation are co-occurrent. We therefore sought to investigate these questions in an iPSC-neuron model. Methods for induction of cortical neurons from iPSCs (iPSC-neurons) through overexpression of the master neuronal transcriptional regulator NGN2 have been well-established to produce excitatory cortical neuronal cells with high speed, consistency, and efficiency^16–18^, and iPSC-neurons are increasingly utilized as a tractable model for neurodegenerative diseases^18–21^. We leveraged these properties to generate direct epigenomic perturbations in a homogenous population of neurons, and to examine downstream effects on gene expression and AD-related pathology.

Here, we performed transcriptomic and functional characterization of iPSC-neurons derived from non-demented control (NDC) donors and from familial Alzheimer’s Disease patient donors harboring an amyloid precursor protein (APP) duplication (APP^Dup^). Our results provide additional evidence for the important role played by chromatin-regulating histone acetylation in neurodegenerative disease, and reveal a homeostatic protective mechanism whereby neurons regulate gene transcription in response to AD pathology.

## Results

### Generation of forebrain-like cortical neurons from APP^Dup^ and NDC iPSCs

We obtained two APP duplication (APP^Dup^) and two non-demented control (NDC) iPSC lines previously established as a model of familial AD^20^. We validated via genomic qPCR that both APP^Dup^ iPSC lines maintained 3 copies of APP compared to 2 copies maintained by both NDC lines (**Supp Fig 1A)**. These cell lines were previously induced using a small-molecule-based neuron induction system and shown to exhibit differences in amyloid and tau species generation upon differentiation^20^. Importantly, the genome and epigenome of these cells have not been explored. Here, we utilized a single-step genetic Ngn2 overexpression strategy to differentiate these lines robustly and consistently to facilitate whole-transcriptome analysis.

**Figure 1.**
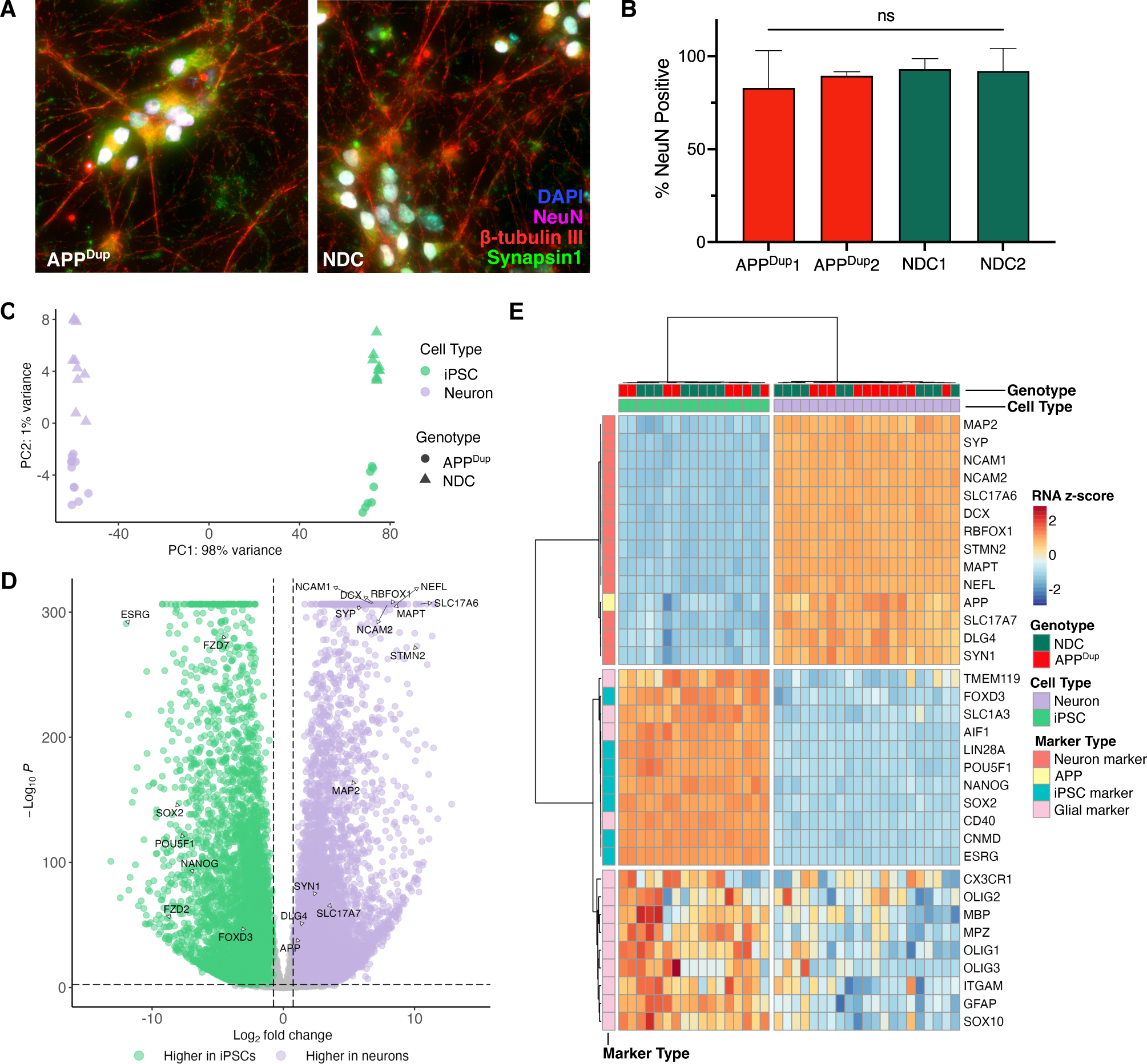
APP^Dup^ and NDC iPSC lines differentiate successfully into excitatory cortical neurons. **A)** Representative immunofluorescence staining for DAPI, beta tubulin III, NeuN and synapsin1 in APP^Dup^ and NDC neurons day 21 CTRL RNAi-treated neurons. **B)** No significant differences in quantification NeuN+ % nuclei were observed among APP^Dup^ and NDC differentiated neurons (*p* = 0.814, one-way ANOVA). Error bars indicate s.e.m. **C)** Principal Component Analysis (PCA) plot of variation of top RNA-Seq genes from differentiated neurons and iPSCs from all cell lines. **D)** Volcano plot depicting differentially expressed genes (DEGs) between APP^Dup^ and NDC CTRL RNAi-treated day 21 neurons and iPSCs. Significance and fold-change threshold of *p-adj.* < 0.005 and log2FC > 0.75 were used. Max -log10(p) value plotted = 300. **E)** Heatmap representing selected neuronal, iPSC, and glial marker gene expression in APP^Dup^ and NDC day 21 neurons and iPSCs.

We introduced cassettes for Tet-On doxycycline-inducible promoter-driven human NGN2, and for EF1α constitutive promoter-driven mApple and blasticidin resistance, into the CLYBL safe harbor locus of each cell line via an established CRISPR-Cas9 transfection strategy^16,17,22^ **(Supp Fig 1B).** Transfected cells were sequentially selected for blasticidin resistance and mApple positivity to establish stable dox-inducible NGN2 transgenic lines. Following established protocols^16,17^, we induced with doxycycline and transitioned to neuronal culture media to direct differentiation of iPSCs into excitatory forebrain-like cortical neurons. Rapid induction of iPSC-neurons through forced overexpression of NGN2 yields functional neurons as measured by electrophysiology recordings by day 21 of neuronal differentiation^16,17,23^, and thus we harvested neurons at day 21 for our experiments. We confirmed mature cortical neuronal identity by immunofluorescence (IF) staining for β-tubulin III, NeuN and Synapsin1 **(Fig 1A)**. No difference was observed in nuclear NeuN positivity between differentiated APP^Dup^ and NDC neurons (**Fig 1B**)

We used RNA-seq to identify transcriptomic changes between iPSCs and 21-day neurons, and consequences of the APP duplication. We observed consistent, broad transcriptional differences between undifferentiated iPSC and differentiated neurons using principal component analysis of differentially expressed genes (DEGs) (**Fig 1C**). We observed 12,496 genes upregulated in differentiated neurons passing a log2FC threshold of >0.75 and p-adj. threshold of <0.005, including neuron associated genes SYN1, MAPT, PSD95 (DLG4), and VGLUT1 (SLC17A7), contrasting with 7398 genes upregulated in iPSCs, including the iPSC-associated pluripotency genes OCT4 (POU5F1), SOX2, ESRG, and NANOG^24,25^ **(Fig 1D**). We used the TissueEnrich R package to examine the most significantly upregulated set of genes following induction into differentiated neurons, and observed that the category with the highest fold-change **(Supp Fig 1C)** and significance **(Supp Fig 1D)** by tissue enrichment were cerebral cortex-associated genes. Additionally, unsupervised hierarchical clustering of selected neuronal, iPSC, and glial marker gene expression in our RNA-seq data revealed robust and distinct clusters made up of differentiated neurons and iPSCs clusters **(Fig 1E)**. We observed robust upregulation of excitatory cortical neuronal marker genes across all differentiated cell lines, and downregulation of iPSC glial marker genes **(Fig 1E)**. As expected, no difference in expression of cell type markers was observed between neurons derived from the APP^Dup^ iPSC and NDC iPSC lines **(Fig 1E)**. Thus, all iPSC lines successfully differentiated into excitatory cortical neurons, and APP duplication status did not alter neuronal differentiation.

### APP^Dup^ and NDC neurons differ in gene expression and Aβ42 secretion

β-amyloid(1-42) (Aβ42) is the dominant peptide found in amyloid plaques in AD patients^3^. Thus, we examined Aβ42 in the AD patient-derived APP^Dup^ neurons using an enzyme-linked immunosorbent assay (ELISA) on neuronal culture media harvested on day 21 of differentiation. Compared to NDC controls, the media from APP^Dup^ neurons contained a significantly higher abundance of Aβ42 (**Fig 2A)**, demonstrating an expected negative consequence of APP duplication.

**Figure 2.**
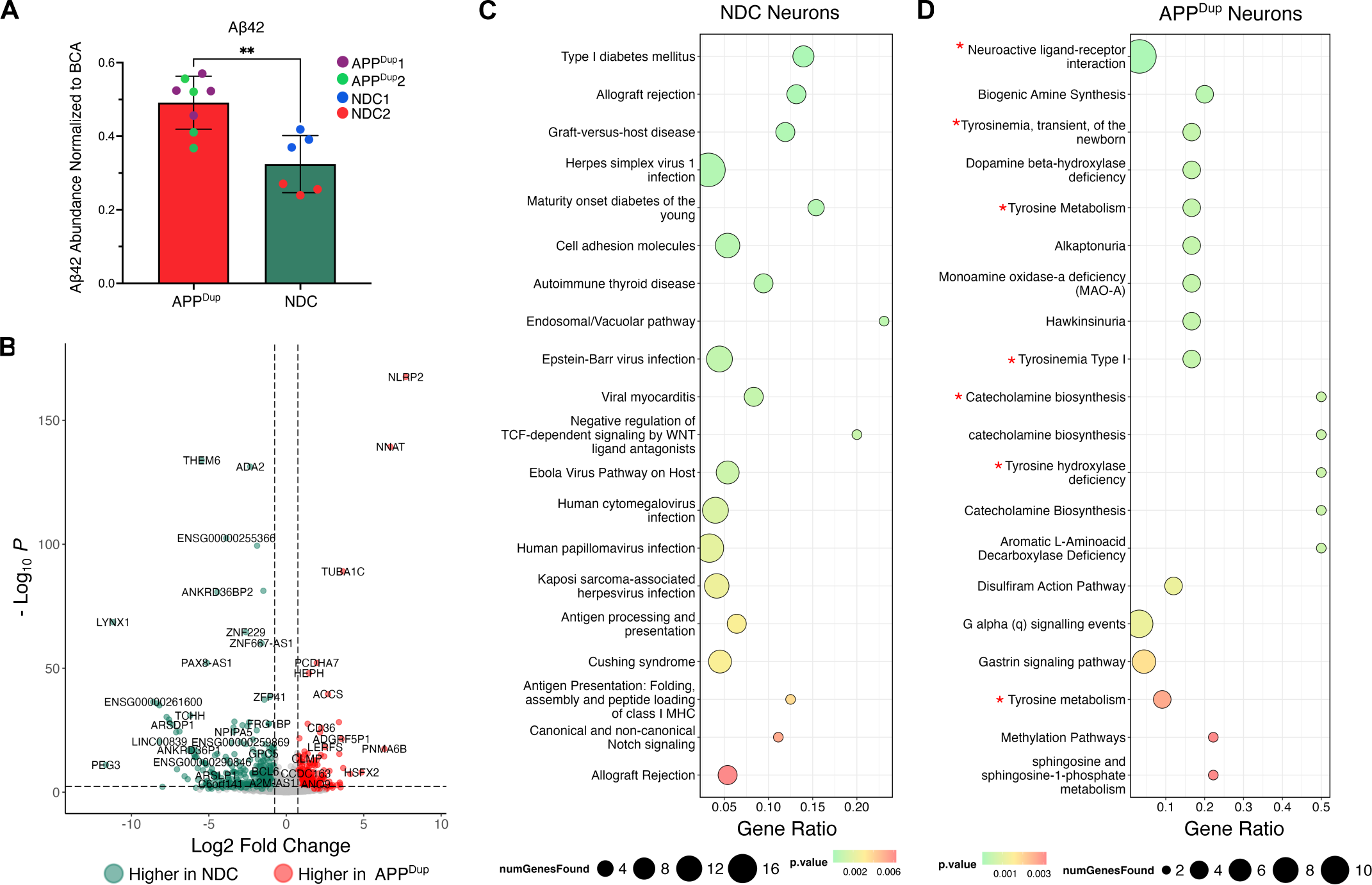
Differences between APP^Dup^ and NDC neurons. **A)** APP^Dup^ day 21 neurons secrete more Aβ42 than NDC day 21 neurons (*p* = 0.0013, two-tailed T-test) at baseline. Media from CTRL RNAi-treated day 21 neurons was collected and measured via enzyme-linked immunosorbent assay (ELISA) and measured Aβ42 abundance was normalized to BCA. **B)** Volcano plot depicting DEGs between APP^Dup^ and NDC day 21 neurons. Significance and fold-change threshold of *p-adj.* < 0.005 and log2FC > 0.75 were used. **C, D)** Bubble plots depicting the top 25 ConsensusPathDB-identified over-represented terms in NDC-neuron- overexpressed **(C)** and APP^Dup^ neuron-overexpressed **(D)** genes. Neuron-related and tyrosine metabolism-related terms are highlighted with an asterisk.

Next, we determined how the transcriptomes of APP^Dup^ derived cells differ from NDC cells at both the iPSC and day 21 neuron stage. We found 1,355 genes upregulated in APP^Dup^ and 593 genes upregulated in NDC iPSCs passing a threshold of log-2-fold-change > 0.75 and *p*-value < 0.005 **(Supp Fig 2A).** At day 21 of differentiation, compared to NDC neurons, 175 genes were upregulated in APP^Dup^ neurons and 332 genes were downregulated (**Fig 2B)**. Plotting the whole-transcriptome gene expression changes in APP^Dup^ vs NDC neurons against the whole-transcriptome gene expression changes in APP^Dup^ vs NDC iPSCs, we observed only a mild correlation of gene expression changes (R = 0.14, **Supp Fig 2B**), establishing that APP^Dup^ affects gene expression differently in iPSCs than in differentiated neurons, and thus the effects of APP^Dup^ on gene expression are context-dependent.

At the iPSC stage, ConsensusPathDB pathway analysis^26,27^ uncovered no significant enrichment of neuron-related terms among significantly upregulated genes in either NDC nor APP^Dup^ cells **(Supp Fig 2C, 2D).** However, neurotransmitter clearance and neuroactive ligand-receptor interaction terms were significantly enriched in APP^Dup^ neurons vs. NDC neurons (**Fig 2C, 2D**). In addition, tyrosine metabolism pathways and monoamine oxidase-a (MAO-A) deficiency pathways were enriched in APP^Dup^ neurons (**Fig 2C, 2D**)—consistent with findings that serum tyrosine is decreased in AD^28^, and that MAO-A catalyzes the cleavage of APP into amyloid species^29^. Importantly, many genes upregulated in APP^Dup^ neurons (red) or downregulated (blue) compared to NDC neurons were identified as involved in the Alzheimer’s Disease Pathway in the Kyoto Encyclopedia of Genes and Genomes (KEGG)^30^ (**Supp Fig 3**). Together, these findings suggest that the underlying baseline transcriptional differences between APP^Dup^ vs. NDC cells are dependent on cell type, and are consistent with known AD-associated pathways.

**Figure 3.**
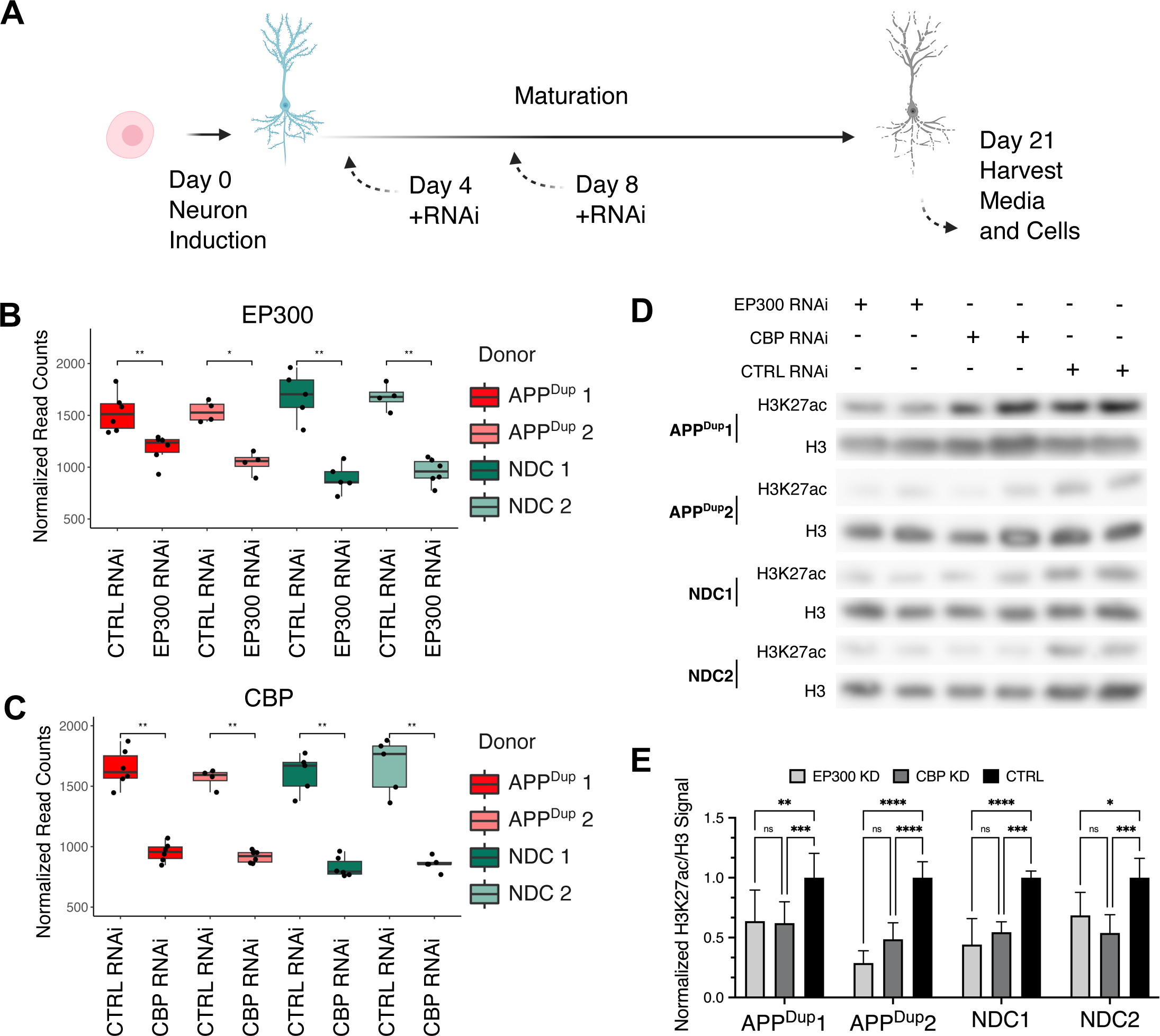
EP300/CBP knockdown leads to reduced H3K27ac in APP^Dup^ and NDC neurons. **A)** Schematic detailing RNAi knockdown during iPSC-neurons differentiation. Cells were transfected with siRNA constructs at Day 4 of neuronal induction and again on Day 8; cells and media for genetic and molecular assays were harvested on day 21. **B, C)** siRNA knockdown of CBP **(B)** and EP300 **(C)** results in reduction of respective transcripts lasting until day 21 (*p* < 0.05 for all comparisons, unpaired two-tailed T-test). Normalized read-counts were obtained by RNA-Seq. **D)** CBP/EP300 KD result in decreased acetylation of H3K27 in both APP^Dup^ and NDC neurons at day 21. Representative Western blots of protein levels of H3K27ac and total H3. **E)** Quantification of H3K27ac/H3 signal across three independent iPSC-neuron differentiations and RNAi treatments.

### EP300/CBP knockdown mediates widespread reduction in gene transcription

In humans, H3K27 is primarily acetylated by the HATs EP300 and CBP^10,31,32^, and is one histone modification of regulatory enhancers and promoters. Therefore, to reduce H3K27ac in our iPSC-neuron model, we employed short interfering RNA (siRNA) to knock down expression of EP300 or CBP in iPSC-neurons (**Fig 3A**). To avoid the potential interference of EP300/CBP depletion in neuronal differentiation, siRNA treatment was initiated on day 4 of the neuronal induction protocol, at which time robust neuronal morphology begins to manifest^16,17^, and then repeated at day 8 (**Fig 3A**).

We analyzed RNAseq levels of EP300 and CBP in treated day 21 neurons, confirming that EP300 and CBP knockdown (KD) was maintained until day 21 of neuronal differentiation (**Fig 3B, 3C**), and corroborated previous reports that RNAi KD persists for weeks in post-mitotic cell types^33^. To determine the effect of EP300 and CBP KD on the overall abundance of H3K27ac, we performed western blot for H3K27ac with overall histone H3 as a loading control. Commensurate with EP300 and CBP transcriptional decrease, H3K27ac was also decreased at day 21 of neuronal differentiation in all cell lines (**Fig 3D, 3E**).

We examined whole-transcriptome gene expression changes via RNAseq following EP300 and CBP KD in APP^Dup^ and NDC day 21 neurons compared to siRNA controls. Upon KD of either HAT enzyme, we observed widespread gene transcription changes in both cell lines that met robustness and significance criteria of log-2-fold-change > 0.50 and *p*-value < 0.005 (**Fig 4A, 4B**). In total, in APP^Dup^ neurons, 1520 genes were downregulated in EP300 (**Fig 4A**) and 1242 genes were downregulated in CBP KD (**Fig 4B)**; in NDC neurons, 1123 and 741 genes were downregulated in EP300 and CBP KD, respectively (**Supp Fig 4A, 4B**). Although EP300/CBP and H3K27ac are primarily associated with enhancer regions and actively transcribed genes^11–13^, we observed mild upregulation of some genes in KD conditions, which may result from secondary effects. Specifically, 768 and 320 genes were upregulated upon EP300 and CBP KD, respectively, in APP^Dup^ neurons (**Fig 4A, 4B**), and 566 and 250 genes were lupregulated upon EP300 and CBP KD, respectively, in NDC neurons (**Supp Fig 4A, 4B**). In summary, we observed anticipated widespread downregulation of gene transcription upon KD of either EP300 or CBP, and the number of affected genes was slightly, but consistently greater in APP^Dup^ neurons compared to NDC neurons.

**Figure 4.**
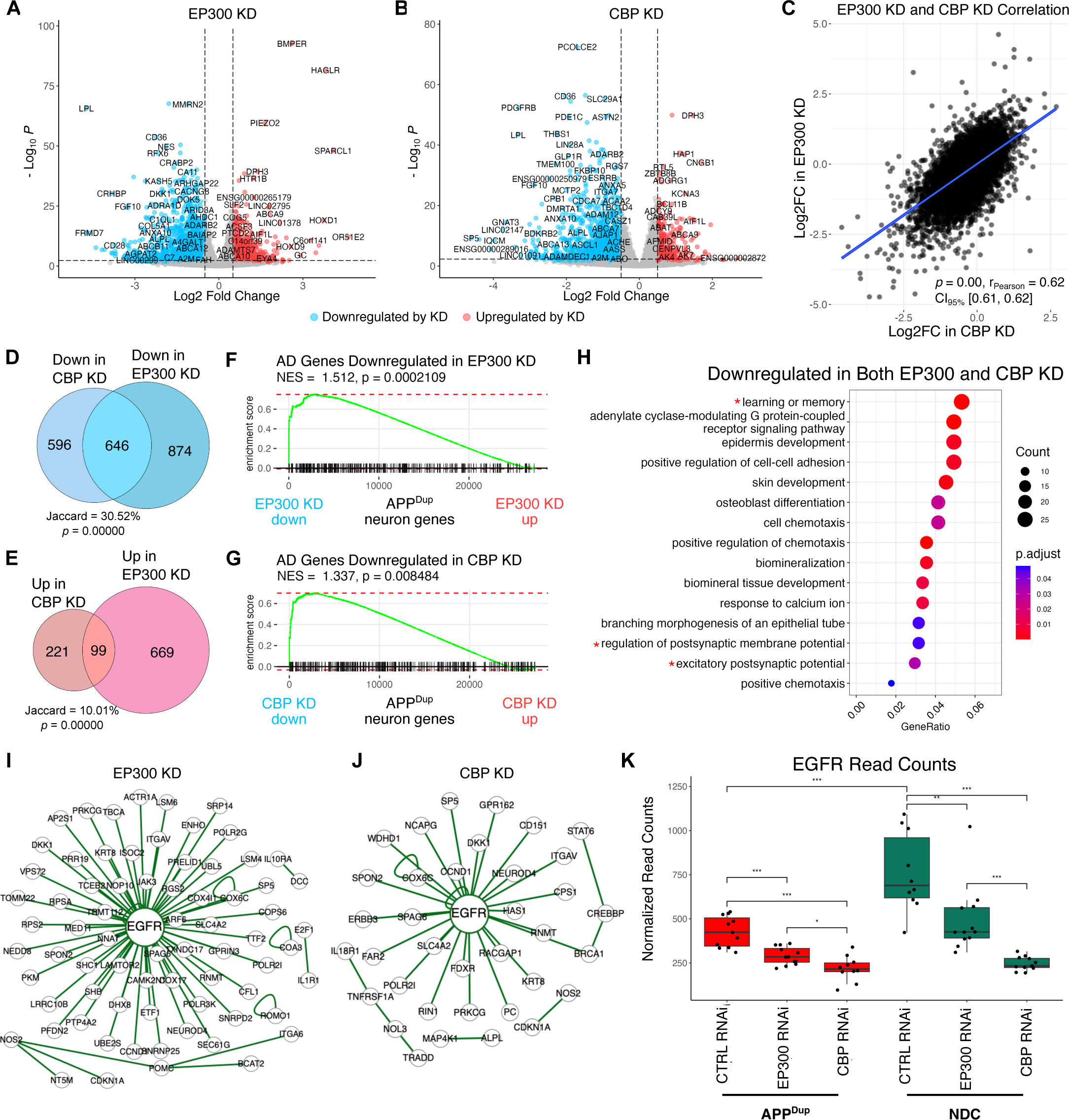
Transcriptional effects of EP300/CBP KD in APP^Dup^ neurons. **A, B)** EP300/CBP KD results in widespread transcriptional downregulation in APP^Dup^ neurons. Volcano plot of gene expression changes between EP300 KD vs CTRL siRNA **(A)** and CBP KD vs CTRL siRNA **(B)** in APP^Dup^ neurons. Significance and fold-change threshold of *p-adj.* < 0.005 and log2FC > 0.50 were used. **C)** EP300 and CBP KD evoke generally similar transcriptional responses. Scatterplot depicting relationship of gene transcriptional changes between EP300 KD and CBP KD (r_pearson_ = 0.62, *p* = 0.00) in APP^Dup^ neurons. **D,E)** Venn diagram showing overlaps of gene expression changes between EP300 and CBP KD in APP^Dup^ neurons. Jaccard index of 412/1355, *p* = 0.00000, hypergeometric test, calculated for overlaps between downregulated genes **(D).** Jaccard index of 32/421, *p =* 0.00000, hypergeometric test, calculated for overlaps between upregulated genes **(E)**. **F, G)** KEGG-identified AD-associated genes are positively transcriptionally controlled by EP300 **(F)** and CBP **(G)** in APP^Dup^ neurons. Gene set enrichment analysis (GSEA) plot depicting placement of KEGG AD genes in transcriptome ranked from highest expressed in CTRL RNAi to highest expressed in EP300 **(F)** and CBP **(E)** KD in APP^Dup^ neurons.**H)** Bubble plot depicting Gene Ontology (GO) terms identified by ClusterProfiler as significantly enriched in genes downregulated in both EP300 and CBP KD in APP^Dup^ neurons. Neuron-related terms are asterisked. **I,J)** EGFR is a central interactor of genes significantly downregulated by EP300 and CBP KD in APP^Dup^ neurons. BioGRID-identified genetic interactions depicted by EasyNetworks (esyN) of genes significantly downregulated by EP300 **(I)** and CBP **(J)** KD. **K)** EGFR transcription is higher in NDC neurons at baseline (+CTRL RNAi) and is decreased by EP300/CBP KD in both APP^Dup^ and NDC neurons (*p* < 0.05 for all comparisons, unpaired two-tailed T-test). Normalized read-counts were obtained by RNA-Seq.

To determine whether EP300 and CBP regulate a similar set of genes, we plotted gene expression changes in EP300 KD vs CBP KD relative to siRNA controls. We observed a strong, significant correlation in whole-transcriptomic RNA expression in EP300 and CBP KD in both APP^Dup^ (**Fig 4C**) and NDC neurons (**Supp Fig 4C**). We examined only the most significantly changed genes in KD conditions in APP^Dup^ neurons: the downregulated genes in EP300 and CBP KD showed a high degree of overlap (Jaccard = 30.5%, p = 0.00000) (**Fig 4D**); in contrast, the upregulated genes showed a much smaller overlap (**Fig 4E**), although the overlap did pass hypergeometric significance testing (Jaccard = 10%; p = 0.00000). A similar degree of overlap was observed in NDC neurons **(Supp Fig 4D, 4E).** Taken together, these data support the view that KD of either EP300 or CBP results in broadly similar effects on overall transcription, with positive gene regulation (i.e. downregulation in enzyme KD) displaying both greater magnitude and similarity between the acetyltransferases compared to negative regulation (i.e. upregulation in enzyme KD).

### Reduction of EP300/CBP affects neuropeptide signaling pathway genes and Alzheimer’s disease pathway genes

We investigated whether EP300 and CBP KD affects genes related to AD, using Gene Set Enrichment Analysis (GSEA) on AD pathway genes identified by KEGG. GSEA revealed that genes downregulated in EP300 and CBP KD were strongly enriched for AD pathway genes in both APP^Dup^ (**Fig 4F, 4G)** and NDC neurons **(Supp Fig 4F, 4G**), indicating that EP300 and CBP drive expression of AD-related genes regardless of APP duplication status. Additionally, KEGG pathway analysis revealed that the majority of the AD-related DEGs in both EP300 (**Supp Fig 5**) and CBP KD (**Supp Fig 6**) in APP^Dup^ neurons are downregulated (blue), supporting the view that EP300 and CBP positively regulate the expression of AD-related genes.

**Figure 5.**
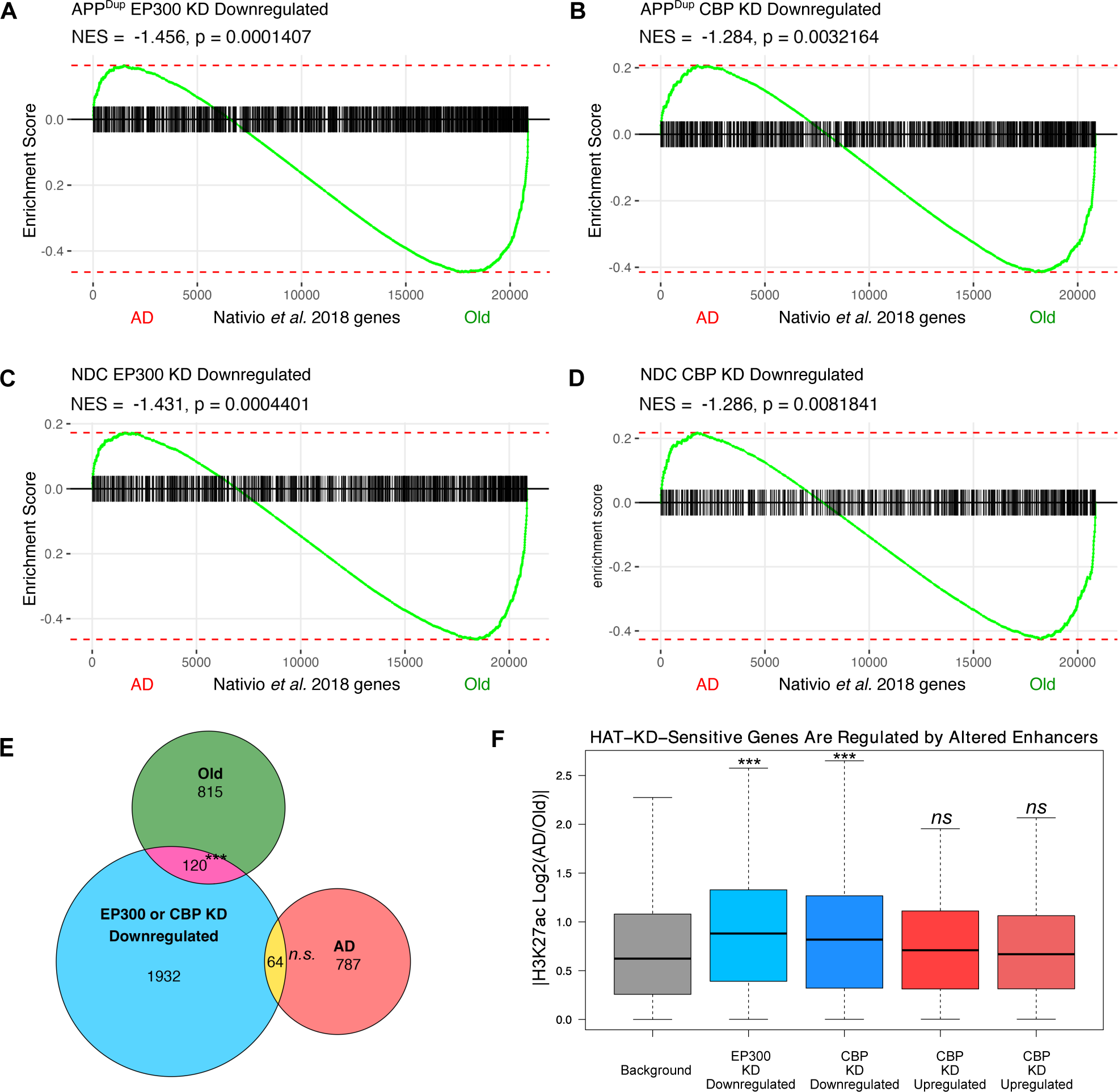
EP300/CBP-regulated genes are affected by AD-associated epigenomic dysregulation. **A-D)** GSEA plots depicting placement of the list of significant EP300-regulated genes in APP^Dup^ neurons **(A)**, CBP-regulated genes in APP^Dup^ neurons **(B)**, EP300-regulated genes in NDC neurons **(C)**, and CBP- regulated genes in NDC neurons **(D)** from this study in the RNA-seq transcriptome of human whole brain, ranked from highest expressed in AD patient brains to highest expressed in non-demented age-matched (“Old”) brains from Nativio *et al. 2018*. **E)** Significant overlap was observed between APP^Dup^ EP300/CBPKD-Downregulated genes and Old-associated genes (N = 120, Jaccard = 120/3718, *p =* 2.61e-08), but not between and APP^Dup^ EP300/CBP KD-downregulated genes and AD-associated genes (N = 64, Jaccard = 64/3718, *p* = 0.613). Hypergeometric statistical testing was performed using the SuperExactTest R package.**F)** EP300- and CBP-regulated genes in APP^Dup^ neurons are significantly associated with either AD-specific or Old-specific H3K27ac enrichment in human postmortem tissue, when contrasted with a control group of HAT-insensitive background genes that have nearby H3K27ac enrichment in the same brains. GREG:STATISTICAL TESTING

**Figure 6.**
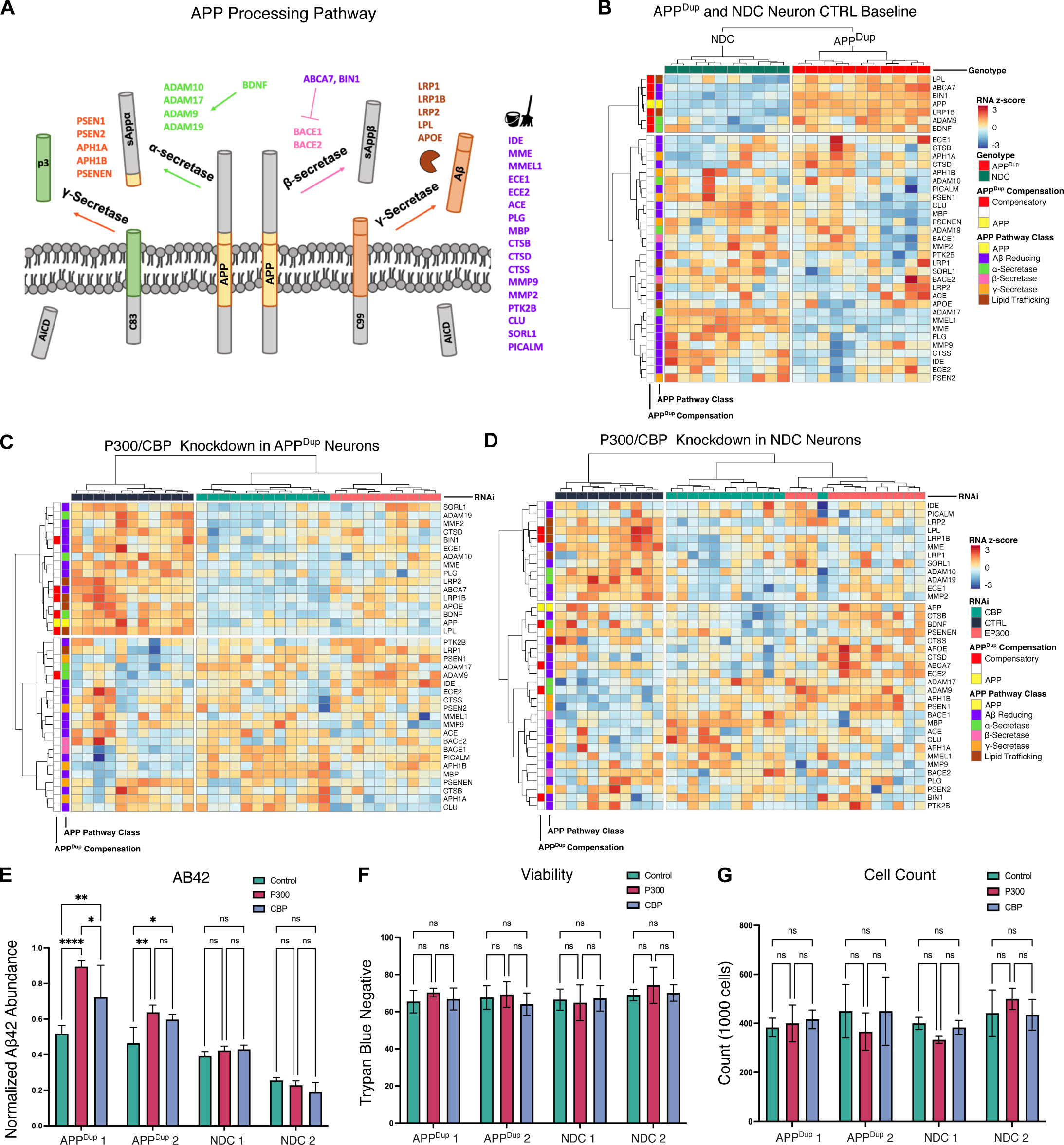
EP300/CBP KD results in downregulation of amyloid-reducing compensatory pathway in APP^Dup^ neurons. **A)** Schematic of analyzed APP processing pathway genes and their functions within the APP processing pathway. **B)** Amyloid-β reduction and α-secretase pathway activation genes are upregulated in APP^Dup^ vs. NDC neurons. Heatmap comparing gene expression of APP processing pathway genes across all APP^Dup^ and NDC CTRL RNAi-treated neurons. **C, D)** Amyloid-reducing genes including those upregulated in APP^Dup^ neurons are downregulated upon EP300/CBP KD in APP^Dup^ neurons **(C)**.Fewer amyloid-reducing genes are downregulated upon EP300/CBP KD in NDC neurons **(D)**. **E)** Aβ42 secretion is increased upon EP300/CBP KD in APP^Dup^ but not NDC neurons. Media from EP300, CBP, and CTRL siRNA-treated day 21 neurons was collected and measured via ELISA and measured Aβ42 abundance was normalized to BCA. **F, G)** EP300/CBP KD does not affect neuron viability or cell count. Day 21 neurons were collected and stained with trypan blue and under counted via hemocytometer microscopy; trypan-blue negativity **(F)** and trypan-blue negative cell count **(G)** were recorded to determine viability and cell count number.

We performed Gene Ontology (GO) analysis^34–36^ on the set of 646 genes downregulated in both EP300 and CBP KD in APP^Dup^ neurons and found that “learning and memory” was the most significantly enriched biological process (BP) GO term with the highest number of genes represented (**Fig 4H**). Also enriched were neuron-related GO terms, such as “regulation of postsynaptic membrane potential” and “excitatory postsynaptic potential” (**Fig 4H**). These findings reflect the known molecular role of EP300 and CBP in learning and memory and neuronal function^37,38^. In NDC neurons, the set of 453 genes downregulated in both EP300 and CBP KD, again “learning and memory” was the most significantly enriched GO term, along with neuron-related GO terms (**Supp Fig 7A**). We performed Cell Compartment (CC) GO analysis on both sets of genes to assess the cellular location of action of EP300- and CBP-controlled genes, and discovered that the neuron projection terminus and axon terminus GO terms were among the highest enriched GO terms in both APP^Dup^ (**Supp Fig 7B**) and NDC neurons (**Supp Fig 7C**). Together, these results confirm that the genes downregulated in EP300 and CBP KD in both APP^Dup^ and NDC neurons are enriched for neuron-related activity.

We performed Easy Networks analysis (eSyn)^39^ to determine genetic associations among the EP300/CBP-regulated gene sets in the biological General Repository for Interaction Datasets (BioGRID)^40^. This analysis uncovered the epidermal growth factor receptor (EGFR) as highly central to the interactions among the genes significantly downregulated in both EP300 KD (**Fig 4I, Supp Fig 7D**) and CBP KD (**Fig 4J, Supp Fig 7E**) in both APP^Dup^ (**Fig 4I, 4J**) and NDC neurons (**Supp Fig 7D, 7E**). EGFR interacts with Aβ42 to synergistically promote memory loss in *Drosophila* AD models ^41,42^, and is increased in an APP/PS1 transgenic AD mouse model ^43^. We examined EGFR expression in EP300/CBP KD, and found that KD in both APP^Dup^ and NDC neurons significantly reduced EGFR expression (**Fig 4K**). We noted lower baseline expression of EGFR in APP^Dup^ neurons compared to NDC neurons (**Fig 4K**), which we discuss below. These findings show central involvement of EGFR in EP300 and CBP KD-mediated gene expression changes in both APP^Dup^ and NDC neurons.

### EP300/CBP-regulated genes are more highly expressed in the transcriptomes of healthy, non-demented aged brains compared to sporadic AD patient brains

In our previous studies of histone acetylation in human postmortem brain, we detected condition-dependent H3K27 acetylation at genomic regions specific to either AD or healthy aging, with a net increase in H3K27ac observed in AD compared to aged normal donors (“Old”)^14^. We also observed a correlation between H3K27ac ChIPseq peak enrichment and RNA-seq gene expression^14^. We therefore investigated here whether the EP300/CBP regulated genes share similarity with gene expression differences between AD patient and Old brains. GSEA revealed that genes downregulated in EP300 and CBP KD (i.e. CBP- and EP300-regulated in our system) were more highly expressed in Old brains in both APP^Dup^ (**Fig 5A, B**) and NDC (**Fig 5C, D**) neurons; we note that a fraction of significant EP300- and CBP-positively regulated genes were highly expressed in AD patients (**Fig 5A- 5D**). We observed a slight but significant overlap between genes downregulated in either EP300 or CBP KD in APP^Dup^ neurons and genes upregulated in Old brain compared to AD brains (Jaccard index = 120/3718 genes, *p* = 2.61e-08), but no significant overlap between EP300/CBP KD-downregulated genes and AD-upregulated genes (Jaccard index = 64/3718, *p* = 0.613) (**Fig 5E**). We observed the same pattern in NDC neurons, where EP300/CBP KD overlapped significantly with Old-upregulated genes (Jaccard index = 80/2226, p =6.65e-06) but not with AD-upregulated genes (Jaccard index = 40/2142, p = 0.75) (**Supp Fig 8A**). Thus, EP300/CBP-regulated genes more closely overlap with a healthy aged transcriptional signature than with a diseased transcriptional signature.

We then examined whether the association between *in vitro* EP300/CBP KD and human brain gene expression is due to the acetyltransferase behavior of these enzymes *in vivo.* To do this, we examined whether genes positively regulated by EP300 and CBP in APP^Dup^ neurons are subject to local, AD-specific depletion of H3K27ac in human brain, as measured by the previously published ChIP-seq datasets. We found that the genes downregulated in CBP/EP300 KD in both APP^Dup^ (**Fig 5F**, blue) NDC neurons (**Supp Fig 8B**, blue bars) were significantly associated with altered H3K27ac enrichment in either AD patient or Old brains; these were compared to upregulated genes (**Fig 5F**, **Supp Fig 8B**, red bars) or background genes unaffected by HAT KD (grey bars). Additionally, the total number of H3K27ac associated peaks per gene in human brain tissue were significantly increased over background for genes downregulated in EP300/CBP KD in both APP^Dup^ and NDC neurons (**Supp Fig 8C, blue bars**); these data suggest that genes regulated by CBP and EP300 in differentiated neurons are subject to more complex regulation in the human brain than expected by chance. In summary, CBP/EP300 regulated genes in APP^Dup^ and NDC neurons are highly expressed in Old brain, and these genes have differential, complex H3K27ac regulation between AD patient and Old brains.

### Reduction of EP300/CBP downregulates amyloid-reducing genes and increase secretion of amyloid-β(1-42)

Given the APP gene duplication, we investigated whether APP pathway-specific genes were differentially expressed between APP^Dup^ and NDC neurons, and if so, whether EP300 and CBP KD alter this. We curated a list of 37 genes involved in APP processing into Aβ in humans (**Fig 6A**). These include genes expected to increase amyloid abundance, such as members of the β-secretase, and γ- secretase complexes. We also included genes expected to contribute to amyloid reduction, either through clearance or decreased production, such as the members of the α-secretase complex and other genes regulating production of amyloid-β species from APP or clearance of amyloid-β (**Fig 6A**).

First, we examined whether expression of the APP-Aβ pathway-associated genes differed between APP^Dup^ and NDC neurons. We performed hierarchical clustering on the normalized RNA-seq expression data in APP^Dup^ and NDC neurons and identified, in addition to APP itself, a cluster of 6 APP pathway-associated genes that were uniformly more highly expressed in APP^Dup^ neurons compared to NDC neurons: *LPL*, *ABCA7*, *BIN1*, *LRP1B*, *ADAM9*, and *BDNF* (**Fig 6B**, top group outlined in red on left). These genes are implicated to reduce amyloid, suggesting that their upregulation in the APP^Dup^ neurons represents a homeostatic, compensatory response to reduce the level of Aβ.

Next, we examined whether EP300 or CBP reduction affects the expression of these and other amyloid pathway genes in APP^Dup^ neurons. We found a cluster of 16 genes downregulated in EP300/CBP KD conditions compared to control siRNA treatment, including 5/6 of the compensatory pathway genes found to be upregulated in APP^Dup^ vs NDC neurons: BIN1, ABCA7, LRP1B, BDNF, and LPL (**Fig 6C**, outlined in red on left). In addition to BDNF, which promotes α-secretase activity and inhibits β- secretase activity^44,45^, the α-secretase complex members ADAM19 and ADAM10 were also decreased upon EP300/CBP KD. The α-secretase complex diverts APP substrate toward α- rather than β- cleavage, reducing Aβ production^46^. ABCA7^47,48^ and BIN1^49–51^ were identified as APP^Dup^ compensatory and also identified as decreased in EP300/CBP KD; these genes reduce amyloid-β by negatively regulating the activity of β-secretase complex member BACE1^47–50^. Additionally, EP300/CBP KD resulted in the decreased transcription of SORL1^52^, MMP2^53^, CTSD^54^, ECE1^55^, MME^56,57^, and PLG^58,59^ which reduce Aβ through amyloid catabolism. We note that some Aβ reducing genes were not downregulated upon KD, however, these genes were more lowly expressed in baseline APP^Dup^ cells (**Fig 6B**). Finally, the lipid trafficking and metabolism proteins APOE, LPL and LRP family members LRP1B and LRP2, which mediate amyloid reduction through co-aggregation with amyloid species and prevention of their extracellular aggregation^21,60–64^, were reduced upon EP300/CBP KD; of these, LPL and LRP1B were also identified as APP^Dup^-compensatory.

Interestingly, the transcriptional response to EP300/CBP KD in NDC neurons differed from APP^Dup^ neurons. Select APP^Dup^-compensatory genes that were decreased by EP300/CBP KD in APP^Dup^ neurons (ABCA7, BIN1, BDNF), were not decreased by EP300/CBP KD in NDC neurons (**Fig 6D**). NDC neurons still showed a cluster of downregulated amyloid-reducing genes upon EP300/CBP KD; however, the total number of affected genes was smaller (12 genes compared to 16 genes in APP^Dup^ neurons), and included only 2/6 of the APP^Dup^ neuron-upregulated compensatory genes: LPL and LRP1B. Hence, genes in the APP-Aβ pathway, particularly those involved in Aβ reduction, were decreasing in EP300/CBP KD, dependent on underlying genetic background of the neurons, with broader downregulation in APP^Dup^ neurons compared to NDC neurons.

While expression of APP itself was increased, as expected, in APP^Dup^ neurons compared to NDC neurons (**Fig 6B**), expression of APP decreased upon EP300/KD KD in both APP^Dup^ and NDC neurons (**Fig 6C, 6D**). It is possible that feedback sensing of a decreased APP abundance induced by HAT KD contributed to the downregulation of amyloid-reducing genes, mirroring the induction of amyloid-reducing compensatory genes in APP^Dup^ neurons. Additionally, the Aβ-producing β-secretase gene BACE1 was significantly upregulated in both EP300 and CBP KD in APP^Dup^ neurons, while CBP KD in NDC neurons resulted in a smaller upregulation of BACE1, and EP300 KD in NDC neurons did not change BACE1 expression (**Supp Fig 9**). These results corroborate our observation that ABCA7, which inhibits BACE1 transcription, decreases upon EP300/CBP KD in APP^Dup^ but not in NDC neurons (**Fig 6C, 6D**). Together, we conclude that the amyloid-exacerbating transcriptional program enacted by EP300 and CBP KD in APP^Dup^ extended to both primary and secondary transcriptional regulation.

We predicted that EP300/CBP KD would increase disease promoting Aβ42 secretion, because (1) Aβ- reducing genes were downregulated in EP300/CBP KD, and (2) EP300/CBP regulated genes were enriched for genes upregulated in healthy aged human brains compared to AD brains. We thus measured, via ELISA, the amount of Aβ42 secreted into the media of APP^Dup^ and NDC day 21 neurons treated with siRNA KD. Treatment with EP300/CBP siRNA compared to control KD increased Aβ42(1- 42) (Aβ42) abundance in both lines of APP^Dup^ neurons, but did not alter Aβ42 abundance in NDC neurons (**Fig 6E**). As controls, there were no differences in cell viability as measured by Trypan blue staining (**Fig 6F**), or in total cell count (**Fig 6G**) resulting from EP300/CBP KD compared to control siRNA in any cell line. Taken together, these data provide evidence that a homeostatic genetic response to increased amyloid-beta production is induced in APP^Dup^ and that the genes involved in this response, along with other amyloid-reducing genes, are regulated by EP300 and CBP.

## Discussion

There exist disparate lines of evidence regarding the nature of epigenomic dysregulation in neurodegenerative Alzheimer’s disease. We and others have reported differential H3K27ac in human brain tissue that is dependent on the brain region surveyed: H3K27ac increases in the lateral temporal lobe^14^, but decreases in the entorhinal cortex of AD patient brains^15^. We also examined the histone acetylation mark H4K16ac and discovered a decrease in the brains of AD patients^9^, and studies in fly models of AD suggest a protective role of H4K16ac against AD pathology-related insults^65,66^ Additionally, in a mouse model of AD, the histone deacetylase HDAC2 increases and H4K12ac decreases^67^, and an ameliorative effect was demonstrated in an AD mouse model of increasing acetyl-CoA synthetase 2 (ACSS2)^68^, which generates acetyl-coA and regulates histone acetylation in rodent hippocampus to promote memory^69,70^. These findings fuel speculation that histone deacetylase inhibitors could be potential therapeutics in Alzheimer’s disease and other neurodegenerative diseases, to overcome a disease-associated “epigenomic blockade”^67,71^.

In this study, we utilized iPSC-neurons derived from familial AD patients with an APP duplication as a model to study the functional effect of reducing the key acetyltransferases EP300 and CBP. KD of the acetyltransferases reduces acetylation at H3K27, and lowers acetylation of other histone and non-histone proteins^10^. In our iPSC-neurons, EP300 or CBP KD reduced total H3K27ac abundance (**Fig 3D, 3E**), which led to widespread downregulation of gene transcription in both APP^Dup^ and NDC neurons **(Fig 4A, 4B, Supp.** Fig. 4A**, 4B)**, including important genes in Alzheimer’s disease pathways (**Fig 4F, 4G, Supp.** Fig. 4F**, 4G, Supp.** Fig 5**, Supp.** Fig. 6). Knockdown of the two enzymes resulted in broadly similar transcriptional changes (**Fig 4C**), and learning and memory and neuron-related processes were affected in both EP300 and CBP KD (**Fig 4H, 4I**). Previous studies of EP300/CBP in the context of AD focused primarily on the well-established roles played by these acetyltransferases in learning and memory^72^. Our study uncovers a novel link between EP300/CBP and amyloid pathology in AD.

Indeed, we find that a subset of genes in the APP-Aβ pathway responsible for amyloid clearance and prevention are strongly upregulated in APP^Dup^ neurons compared to NDC neurons (**Fig 6B**), suggesting activation of a homeostatic genetic response to compensate for increased abundance of APP and production of downstream Aβ species. Compared to knockdown in NDC neurons, knockdown of either EP300 or CBP in APP^Dup^ neurons resulted in stronger downregulation of genes with amyloid-reducing function (**Fig 6C, D**), including the APP^Dup^-upregulated, compensatory genes mentioned above.

Importantly, likely as a consequence of downregulation of these amyloid-reducing gene programs, knockdown of EP300 or CBP causes an increase in abundance of disease-associated Aβ42 secreted by APP^Dup^ neurons **(Fig 6E)**.

Our findings reveal several pathways potentially impacted by EP300/CBP KD leading to increased toxic Aβ42. LPL is strongly expressed by neurons^73^ and is involved in extracellular association and sequestration of amyloid-β. Interestingly, we discovered that LPL was among the most significantly downregulated genes in neurons with EP300/CBP KD **(Fig 4A, 4B).** Consistent with this finding, a small molecule inhibiting EP300/CBP catalytic activity strongly decreases LPL expression in mouse adipocytes^74^. LPL is an amyloid-β-binding protein promoting cellular uptake and clearance of amyloid-β in astrocytes^60^. Taken together, downregulation of LPL by KD of EP300/CBP may lead to increased amyloid-β. Although glial cells are considered to be primarily responsible for uptake and clearance of amyloid-β^21,75^, our results open the possibility that LPL may have glial cell-independent effects on amyloid reduction, warranting further analysis. Decreased expression of BDNF may have contributed to the increase in Aβ in EP300/CBP KD cells. BDNF reduces Aβ production^46^ by promoting α-secretase activity to the exclusion of β-secretase activity^44,45^ We find that EP300/CBP knockdown reduces BDNF expression, suggesting it could be a candidate for amyloid-targeting therapeutic approaches.

While our transcriptional and phenotypic data suggest a physiologic amyloid-reducing role of EP300/CBP, we did identify EGFR as a downstream central interactor of both EP300- and CBP- controlled gene expression (**Fig 4I, 4J**), and showed that both EP300 and CBP KD reduced EGFR expression in APP^Dup^ and NDC neurons (**Fig 4K**). EGFR has been implicated in mediating neurotoxicity downstream of Aβ, primarily via utilization of EGFR inhibitors, which reduce amyloid-associated inflammation and ameliorate memory resulting from introduced amyloid^41,76–78^. Upregulation of the EGFR ligand, epidermal growth factor (EGF), reduces amyloid-related deficits without affecting levels of Aβ itself^79^. Therefore, it is likely that EGFR-mediated neurotoxicity is dependent on but does not contribute to amyloid pathology, and occurs after Aβ accumulation has already been established, falling downstream of the scope of our study. Nevertheless, these results indicate that although EP300/CBP regulate a protective amyloid-reducing pathway, they concurrently regulate EGFR, an Aβ toxicity- exacerbating interactor, underlining that there is a complex relationship between histone acetylation, gene activation, and the ultimate phenotype of Aβ pathology and neurotoxicity.

The limitations of our study reflect the limitations inherent in our model system. While an established single gene mutation, like APP duplication in the familial AD lines used here, likely leads to a larger and more consistent effect size on measurements compared to the use of sporadic AD lines, the low number of cell lines used in our study poses a valid concern for generalizability. Additionally, our model lacks astrocytes or other glial cell types, which may explain discrepancies between our study and others conducted in heterogenous brain tissue. For example, while we observed increased AD pathology upon KD of EP300/CBP in the form of Aβ42 secretion (**Fig 6E**), it is possible that the exclusively neuronal makeup of our model prevented us from observing potential beneficial effects of EP300/CBP KD. such as the downregulation of noxious agent EGFR, which appears to enact its Aβ42- worsening effect through neuroinflammation and its interaction with glial cells^77,78^. Of particular relevance, our previous study reporting a net increase in H3K27ac in AD brains was performed in whole brain tissue, which does indeed contain glial cell types. Thus, it is possible that different cell types experience different degrees of histone acetylation change; for example, a report examining histone acetylation changes in individual cell types identified oligodendrocytes as the primary cell type to experience H3K27ac peak increases associated with amyloid load^80^.

Additionally, we limited the scope of our experiments to the effect of EP300 and CBP KD on H3K27ac and amyloid pathology. We did not investigate the relationship of these mechanisms to tau, though tau acetylation has been shown to contribute to tau pathology and dementia^81,82^. Whether similar compensatory mechanisms against tau pathology progression exist in tau mutant iPSC-neuron models, and whether EP300/CBP and H3K27ac drives the expression of those homeostatic programs in those models, remain interesting potential future directions. Additionally, while we demonstrated that neither CBP nor EP300 KD impacted neuronal viability (**Fig 6F**), we did not investigate in detail the potential loss of normal neuron function, an especially important consideration, given the well-established role of EP300 and CBP in learning and memory^37,38,83^.

In summary, our findings suggest a complex mechanism of disease-associated EP300/CBP dysregulation and provide further evidence of the crucial role EP300/CBP plays in AD-related pathological processes. In contrast to postmortem brain tissue, in which functional experiments are not feasible, utilization of an iPSC-neuron *in vitro* model allowed us to address, via perturbation experiments, the role HATs play in AD. In particular, using iPSCs derived from familial AD patients with an APP duplication allowed us to isolate the role EP300/CBP plays in Aβ pathology. We show that, inneurons, EP300/CBP KD lowers H3K27ac, inhibits the expression of genetic programs compensating for increased Aβ load, and leads to increased amyloid-β secretion. Future strategies targeting the reduction or increase of histone acetyltransferases as a potential therapeutic mechanism should thus consider the broad role played by H3K27ac as a general activator of transcription and cell-type dependent effects.

## Materials and Methods

### iPSC Lines and Maintenance

APP^Dup^ and NDC iPSC lines were previously characterized and reported^20^ and were the generous gift of the lab of Lawrence Goldstein, University of California, San Diego (lines APPDp1.1, APPDp2.1, NDC1.1, NDC2.1). Established practices for maintenance and passaging of iPSC cell lines were followed.^17,20,84^ Briefly, freshly passaged undifferentiated cells were plated on hESC-qualified Matrigel (Corning 354277)-coated tissue-culture treated 6-well plates in Essential 8 media (Gibco A1517001) containing 10 uM ROCK inhibitor Y-27632 (Tocris 1254). Media without ROCK inhibitor was exchanged daily and wells were passaged at approximately 70-80% confluence by washing with DPBS and mechanically dissociating with ReLeSR (StemCell 100-0483) into cell cluster suspensions. Suspended cell clusters were frozen in Stem Cell Freezing Media (ACS-3020). Cells were routinely checked for mycoplasma contamination.

### Verification of APP copy number

Verification of APP copy number was performed by quantitative PCR (qPCR) of two exon-intron junctions of APP on genomic DNA of all cell lines, normalized to genomic β-globin (*HBB*). The following primers were used for qPCR analysis: tcttcctcccacagctcctggg (FW) and gcattagccacaccagccacca (RV) (*HBB*); ttcccacccttaggctgctggt (FW) and agccttcaccttagggttgccca (RV) (*HBB*); gccaacgagagacagcagctgg (FW) and aactcggctgcagcgagaccta (RV) (*APP*); aaccaccgtggagctccttccc (FW) and ccttgctggctcaggggactct (RV) (*APP*).

### Transfection of Cell Lines with NGN2 Construct

Creation of doxycycline-inducible NGN2 cell lines was performed on all fAD and NDC cell lines according to a previously described method.^17^.All plasmids used were the generous gift of Michael Ward. Briefly, plasmid with blasticidin resistance cassette and mApple were transfected into cells. Cells were passaged and treated with by puromycin and sorted by fluorescence-activated cell sorting (FACS) for mApple positivity after 2 passages with blasticidin to select for cells with long-term stable insertion of the transgene cassette.

### iPSC-Neuron Differentiation

Differentiation of iPSC-neurons was based an established protocol for single-step NGN2-mediated induction of cortical neurons with minor modifications.^17^ Briefly, 80% confluent wells of undifferentiated iPSCs were mechanically dissociated to single cells with StemPro Accutase (Gibco A1110501), then plated at a density of 500k cells/well in a Matrigel-coated 6 well-plate with ROCK inhibitor (day -1).

Media was changed to Neuron Induction Media 24 hours after plating (day 0) and cortical neuron differentiation was induced with doxycycline (Sigma D9891). Fresh Neuron Induction Media containing doxycycline was exchanged on day 1. On day 2, wells were washed with DPBS, mechanically dissociated to single cell suspension with accutase, and plated at a density of 1 million cells per 6-well plate (or 375k cells per 12-well plate) in poly-L-ornithine (Sigma P3655) -coated wells in Neuron Differentiation Media with 10 uM ROCK inhibitor. Cells were treated for 24 hours with 5 uM araC (Tocris 45-205-0) and media was replaced on day 3 with fresh Neuron Differentiation Media without ROCK inhibitor or araC. On day 4, media was replaced with Neuronal Media, and beginning on day 8, half-volume media changes with Neuronal Media was performed every 4 days until harvest. Media was collected for ELISA assays and cells were harvested with mechanical dissociation alone for RNA-seq and Western blotting assays. Cell viability was determined via Trypan Blue (ThermoFisher) staining and counting on a hemocytometer under a bright field microscope.

### RNAi knockdown / Transfection

siRNA duplexes against CBP, EP300, and a scrambled control were designed and synthesized by Horizon Discovery Biosciences Ltd. On Day 4, siRNA duplexes were diluted in Opt-MEM (Gibco 31985062) and combined in Lipofectamine RNAiMax transfection reagent (ThermoFisher 13778150) according to the manufacturer’s instructions. Mixture was added to Neuron Media at a final siRNA concentration of 25nM and media exchange on day 4 was performed as described above. On day 8, transfection was repeated with an additional dose of Opti-MEM, Lipofectamine RNAiMax, and Neuronal Media, and siRNA duplex at a final concentration of 50 nM, concurrent with a half-volume media change, for a total of two RNAi KD treatments spaced 4 days apart on Day 4 and Day 8 respectively.

### RNA-Seq

Frozen cell pellets from at least two independent rounds of differentiation were suspended in TriZol reagent and homogenized with handheld tissue homogenizer and extracted as previously described.^85^ Trizol-extracted RNA samples were quantified on Qubit before cDNA synthesis and library prep with NEB kit, then quantified with NEB Library Quant Kit before sequencing on Illumina NextSeq 550 platform. RNA-seq tag reads were aligned to the GRCh38/hg38 reference human genome assembly using STAR with default parameters.

### Immunofluorescence

For immunofluorescence experiments, induction of iPSCs into cortical neurons was performed as described above with the modification of plating cells on poly-L-ornithine coated glass coverslips on Day 2. At indicated neuronal maturity time points, immunofluorescence and imaging was performed as described previously.^86^ Briefly, cells were fixed in 4% paraformaldehyde in PBS for 10 minutes at room temperature, washed in PBS, permeabilized in 0.5% Triton-X solution for 10 minutes, and then blocked at room temperature in 10% BSA-PBS solution. Fixed and permeabilized cells were then incubated with primary antibody in 5% BSA-PBS + 0.1% Tween 20 solution at 4 °C overnight. Incubation with Alexa Fluor-coupled secondary antibody was performed at room temperature for 1 hour without shaking and washed in PBS. Slides were mounted in VECTASHIELD Antifade mounting medium with DAPI (Vector Laboratories H-1200-10) and imaged on a Nikon Eclipse Ti2. Images were analyzed in Fiji.

### Western blotting

Frozen cell pellets from three independent rounds of differentiation were lysed in Tris buffered saline (TBS) containing 1% NP-40 and 1 mM freshly prepared MgCl_2_ and 1:100 Halt Protease inhibitor cocktail (ThermoFisher 78430). SDS was added to a final concentration of 1% and samples were boiled at 95°C for 10 minutes. Samples were spun at top speed in a tabletop centrifuge and quantified by Qubit and BCA protein quantification assay before loading in a NuPage 4-12% Bis-Tris gel (ThermoFisher NP0322BOX) and subjected to electrophoresis. Samples were transferred onto a 0.2 μm PVDF membrane and blocked for one hour at room temperature in 5% milk-TBS solution containing 1% Tween (TBST). Primary antibodies were added in 5% milk-TBST solution overnight, followed by three washes in TBST. Secondary antibody conjugated to HRP was performed in 5% TBST for 2 hours before 3 washes in TBST. Membrane was imaged by Fujifilm LAS-4000 imager. Quantification of blot intensity was performed in ImageStudioLite v5.2.5. Ratio of H3K27ac intensity to H3 intensity was taken and normalized to CTRL RNAi H3K27ac/H3 intensity for each batch to control for batch variation in intensity. Bar plot and statistical testing (Tukey’s multiple comparison test) were prepared in Prism 10 software v10.0.0.

### Data analysis

For all analyses, raw counts files were loaded into R and quality control metrics of total read counts >7500 for each sample and > 5 read counts in at least 3 samples were applied. For RNAi treatment vs CTRL RNAi comparisons, design = ∼treatment +genotype+batch was used to model the effect of treatment and genotype on transcription, with comparisons being specified between RNAi treatment and CTRL RNAi downstream of linear model building, and a log2 fold-change cutoff of >0.50 was chosen. For APP^Dup^ vs NDC comparisons, design = ∼genotype+batch was used to model the effect of genotype on transcription, the same adj.-p-value cutoff of 0.005 was chosen, but a relatively stringent log2 fold-change cutoff of >0.75 was chosen to adjust for the large number of identified DEGs. DEG identification and PCA plot generation from the default parameter of N = top 500 differentially expressed genes in each comparison was performed using the DESeq2 R package version 1.38.3^87^. Gene expression was normalized for library size and dispersion through the median ratio method provided through the estimateSizeFactors function of DESeq2 and corrected for both batch and donor variation through use of the limma package version 3.54.2^88^. Visualization of KEGG AD pathway involvement of gene sets was generated using the pathview package v1.38.0^89^. Tissue enrichment analysis was performed using the TissueEnrich package v1.18.0.^90^ Differentially regulated genes were visualized using the enhancedVolcano^91^ package v1.16.0. Heatmaps were generated by the pheatmap package developed by Ravio Kolde (University of Tartu) v1.0.12. Dendrograms drawn and arranged using the Ward.D agglomeration method and dendrogram distances were determined by Pearson correlation distance within the pheatmap package. Differentially expressed genes between iPSC and differentiated neurons were analyzed using ConsensusPathDB Release 35 (05.06.2021)^27^. Gene expression enrichment was calculated using the fGSEA^92^ R package v1.24.0 based on the Gene Set Enrichment Analysis (GSEA) algorithm^93,94^. ClusterProfiler^36^ v4.6.2 was used to perform Gene Ontology analysis of differentially expressed genes in EP300/CBP KD. Box plots were generated using ggplot2^95^ R package v3.4.3. All analyses described above were performed with a random seed of 865 for reproducibility. The esyN v2.1 Graphs tool^39^ was used to draw gene network plots, relying on BioGRID v4.4.225 and using high- and low-throughput genetic (not physical) interactions. For all analysis above, R v3.16 was employed. Additionally, Python v3.10.8 was used to process the Nat Genet 2020 Nativio et al RNA-seq and chIP-seq data and create the boxplot in 5F, filtering out genes without H3K27ac peaks and without gene expression measurements. Where multiple peaks targeted a gene, the peak with the largest chIP- seq alteration score (|log2(H3K27ac AD/Old)|) was chosen as the representative peak for that gene, lending its score for the boxplot. The background gene list was chosen to include none of the genes in any of the CBP or EP300 KD differential gene lists, randomly sampling from all other genes in the filtered list to produce a new list with a number of genes in the largest such DEGs list (the random sample seed for this analysis was set to 100 for reproducibility).

### Statistics

Statistics for Aβ42 ELISA, NeuN positivity, cell count, and viability were performed within GraphPad Prism 10 software. All data are presented as mean + s.e.m. One-way analysis of variance (ANOVA or one-tailed Student’s *t-*tests were performed unless otherwise noted. Statistical significance was set to *p* < 0.05 for all experiments except for DESeq analyses of in vitro neuron RNA-seq data, for which significance threshold was set more stringently, *p*-adj. < 0.005. Statistics for analyses of high-throughput data were performed within R with the ggplotstats (https://indrajeetpatil.github.io/ggstatsplot/) and ggsignif^96^ R packages. Hypergeometric tests for overlap between gene sets were performed using the SuperExactTest R package^97^, with the universe of potential overlapped genes set to the intersection of genes with mapped reads in each experiment performed. For Figure 5F, Supp Fig 8B, Supp FIg8C, permutation test was performed with the coin library v1.4.2 in R.

## Supporting information

Supplemental Figures with Legends

