## Supplemental Figures with Legends for "Histone acetylation in an Alzheimer’s disease cell model promotes homeostatic amyloid-reducing pathways"

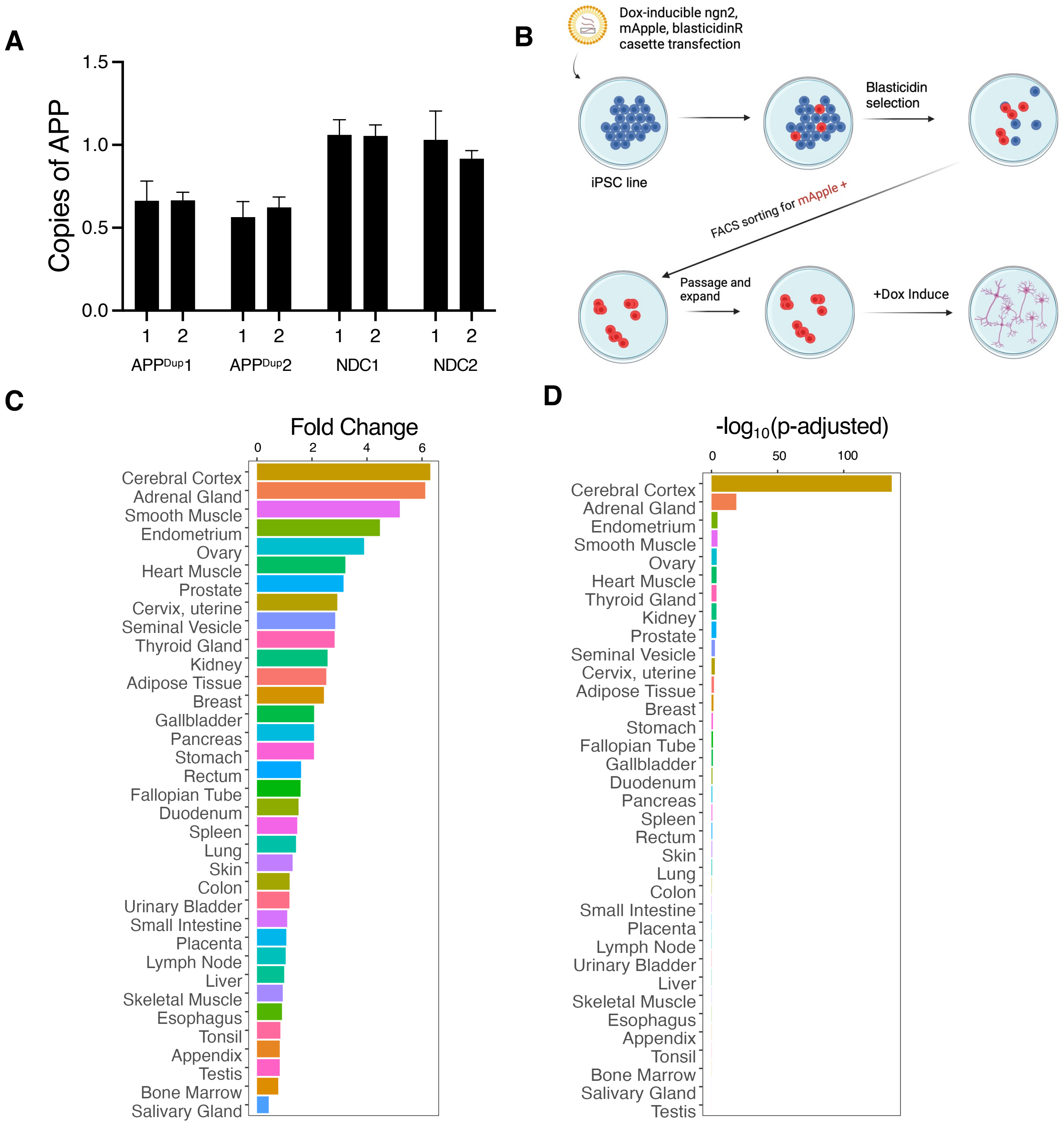


**Supplementary Figure 1. APP$^{Dup}$ and NDC neurons exhibit robust excitatory cortical neuron identity after single-step NGN2 induction. A)** Quantitative PCR (qPCR) of two intron-exon junctions in *APP* locus performed on genomic DNA isolated from from APP$^{Dup}$ and NDC iPSCs, normalized to genomic beta-globin levels. **B)** Schematic of general workflow of ngn2-inducible iPSC line development. **C, D)** Differentiated APP$^{Dup}$ and NDC neurons express genes associated with cerebral cortical tissue. Cerebral cortex-associated genes are the most highly enriched gene set expressed in differentiated neurons compared to iPSCs, ranked by both fold-change **(C)** and significance **(D)**.

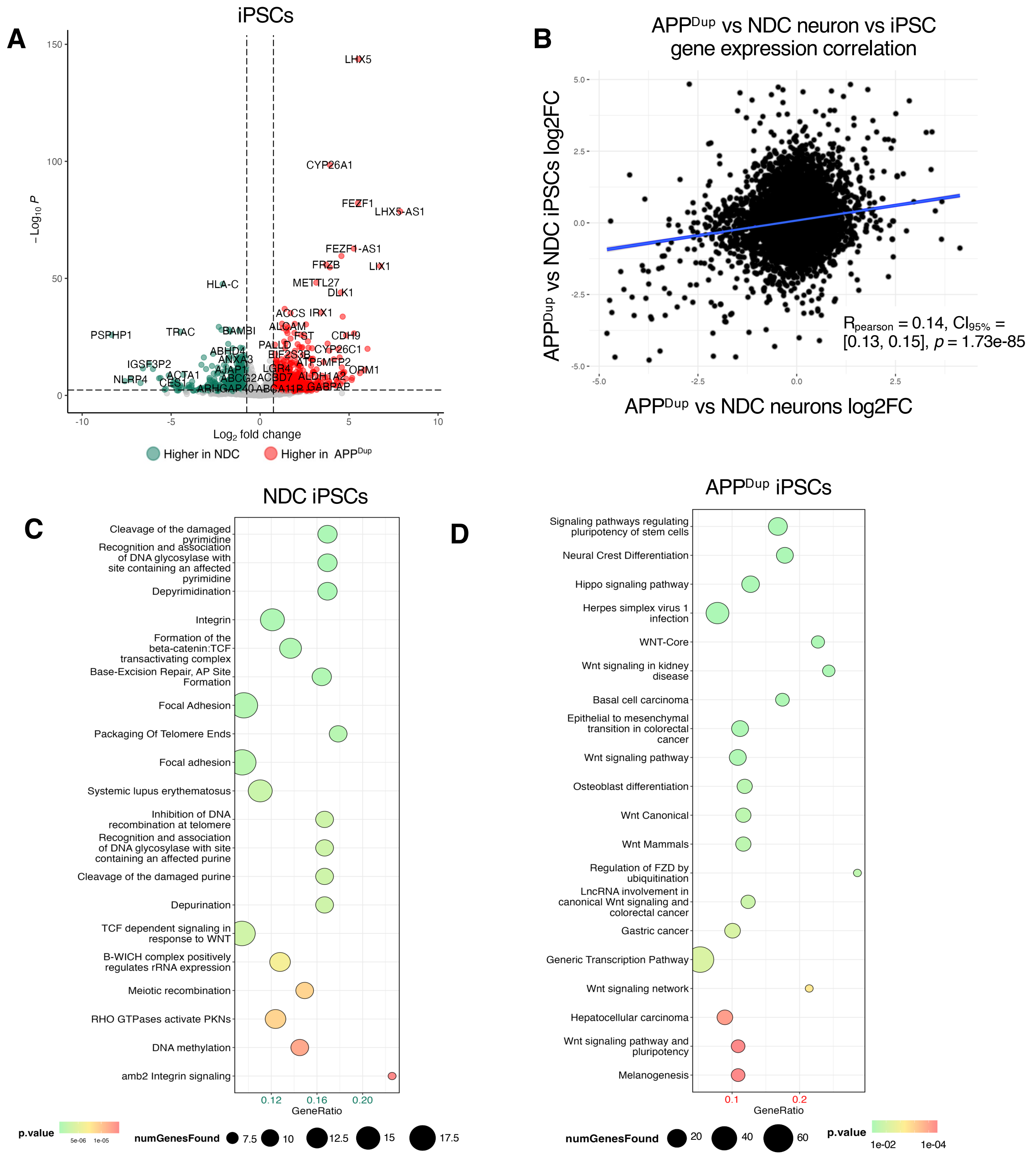


**Supplementary Figure 2. Transcriptional differences between APP$^{Dup}$ and NDC iPSCs. A)** Volcano plot showing baseline gene expression differences between APP$^{dup}$ and NDC iPSCs. **B)** Scatterplot showing correlation between transcriptional gene expression fold-change between APP$^{Dup}$ vs NDC neurons and APP$^{Dup}$ vs NDC iPSCs ($r_{Pearson} = 0.14$, $p = 1.73e\text{-}85$). **C, D)** Bubble plots depicting top 25 terms identified by ConsensusPathDB as overrepresented in NDC **(C)** and APP$^{Dup}$ **(D)** iPSCs.

#### APP$^{Dup}$ vs NDC Neuron KEGGview

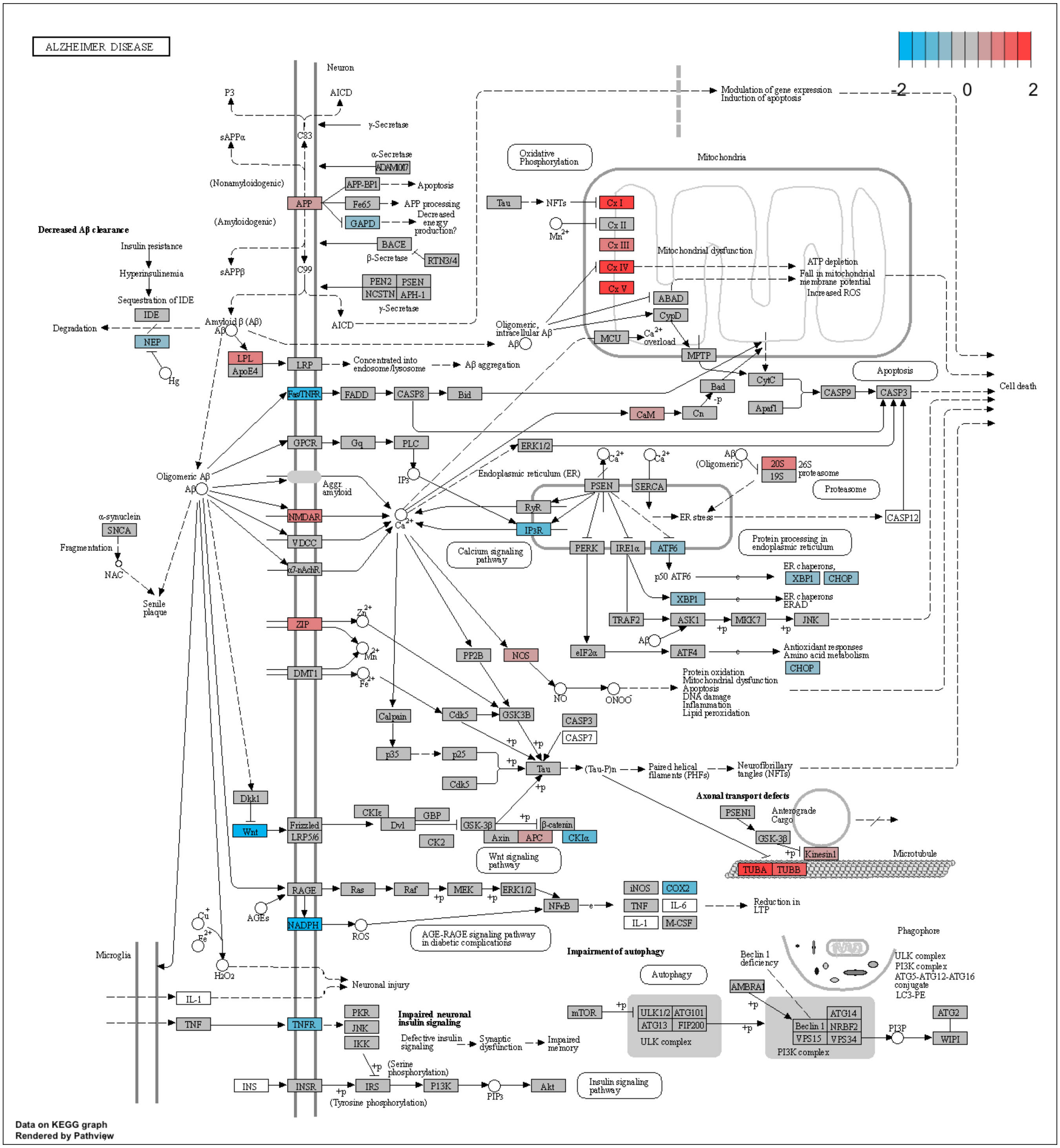


**Supplementary Figure 3. AD Pathway Gene Expression is Affected by APP duplication in neurons.** KEGG pathway view of AD-related differential gene expression pattern between APP$^{Dup}$ and NDC neurons generated in Pathview. Genes are color-coded based on expression from highest in APPDup (red) to highest in NDC (blue).

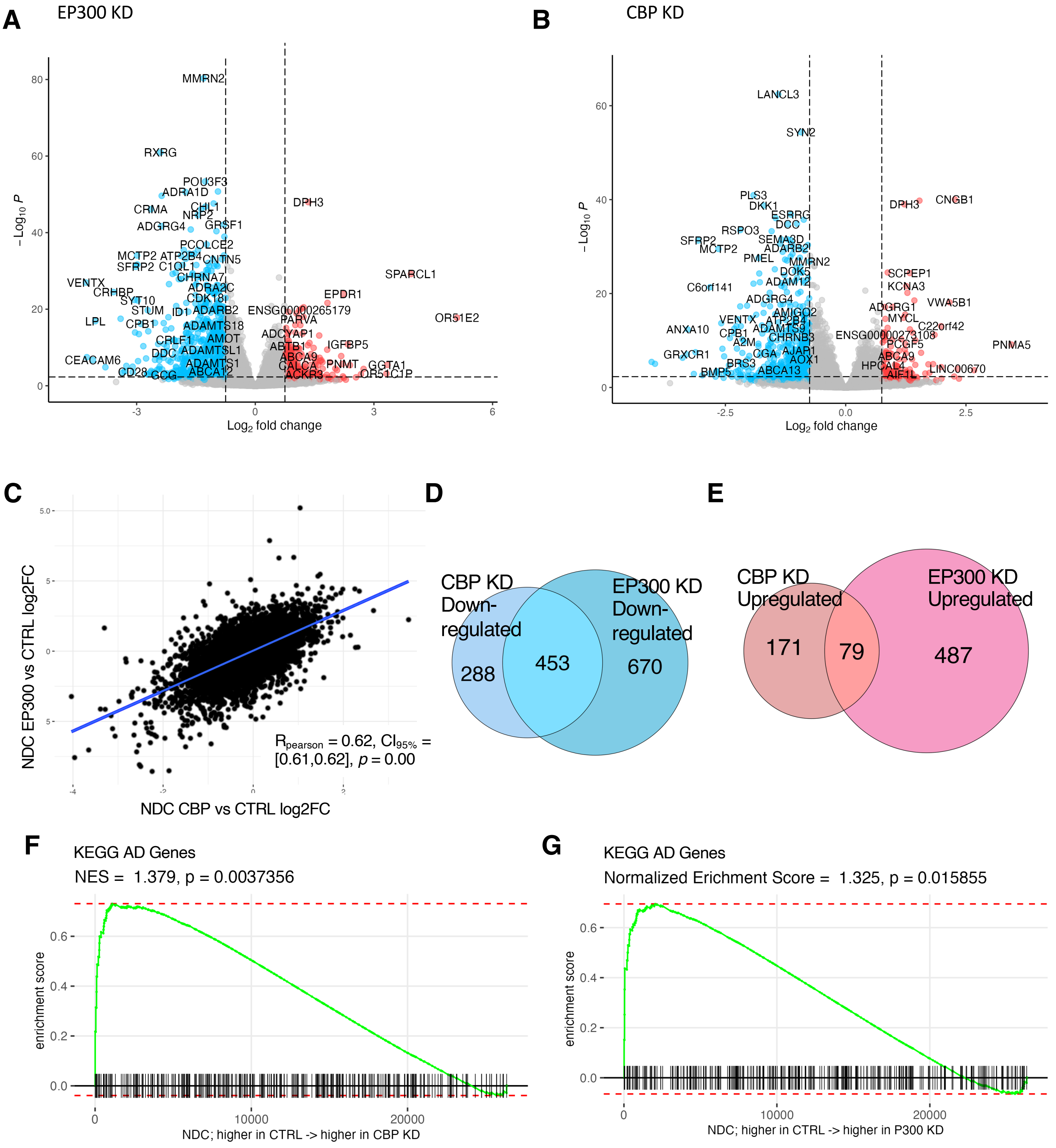


**Supplementary Figure 4. Transcriptional effects of EP300/CBP KD in NDC neurons. A, B)** EP300/CBP KD results in widespread transcriptional downregulation in NDC neurons. Volcano plot of gene expression changes between EP300 KD vs CTRL siRNA **(A)** and CBP KD vs CTRL siRNA **(B)** in NDC neurons. **C)** EP300 and CBP KD evoke generally similar transcriptional responses. Scatterplot depicting relationship of gene transcriptional changes between EP300 KD and CBP KD ($r_{pearson}$ = 0.62, $p$ = 0.00) in NDC neurons. **D,E)** Venn diagram showing overlaps of gene expression changes between P300 and CBP KD in NDC neurons. Jaccard index of 453/1411, $p < 0.05$, hypergeometric test, calculated for overlaps between downregulated genes **(D).** Jaccard index of 79/737, $p$ = 0.05, hypergeometric test, calculated for overlaps between upregulated genes **(E)**. **F, G)** KEGG-identified AD-associated genes are positively transcriptionally controlled by EP300 **(F)** and CBP **(G)** in NDC neurons. Gene set enrichment analysis (GSEA) plot depicting placement of KEGG AD genes in transcriptome ranked from highest expressed in CTRL RNAi to highest expressed in EP300 **(F)** and CBP **(G)** KD in NDC neurons.

#### EP300 KD vs CTRL APP$^{Dup}$ Neurons

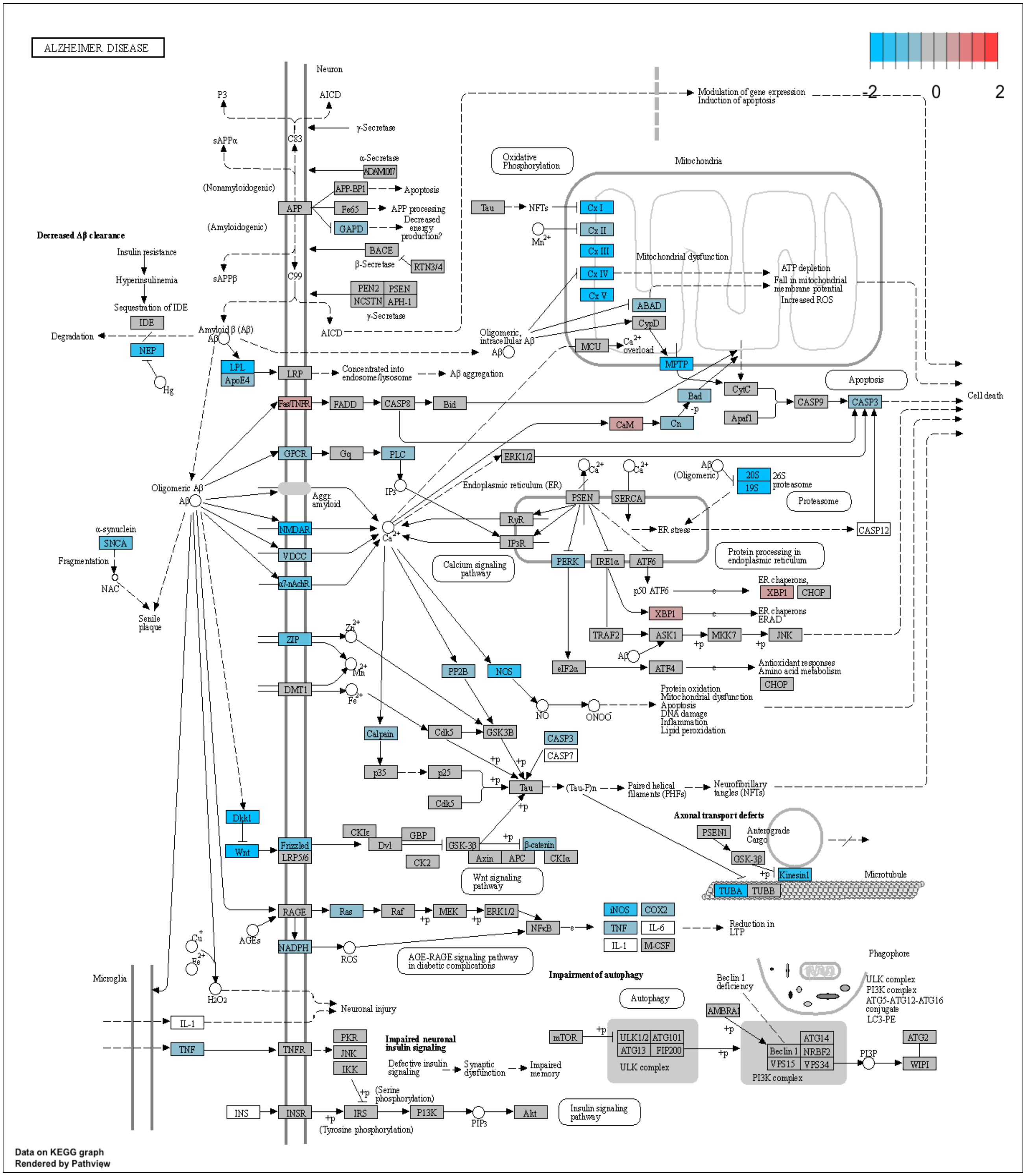


**Supplementary Figure 5. EP300 KD Results in Widespread Downregulation of AD Pathway genes.** KEGG pathway view of AD-related differential gene expression pattern between and EP300 KD and CTRL RNAi in APP$^{Dup}$ neurons generated in Pathview. Genes are color-coded based on expression from highest in KD (red) to lowest in KD (blue).

**CBP KD vs CTRL APP$^{Dup}$ Neurons**

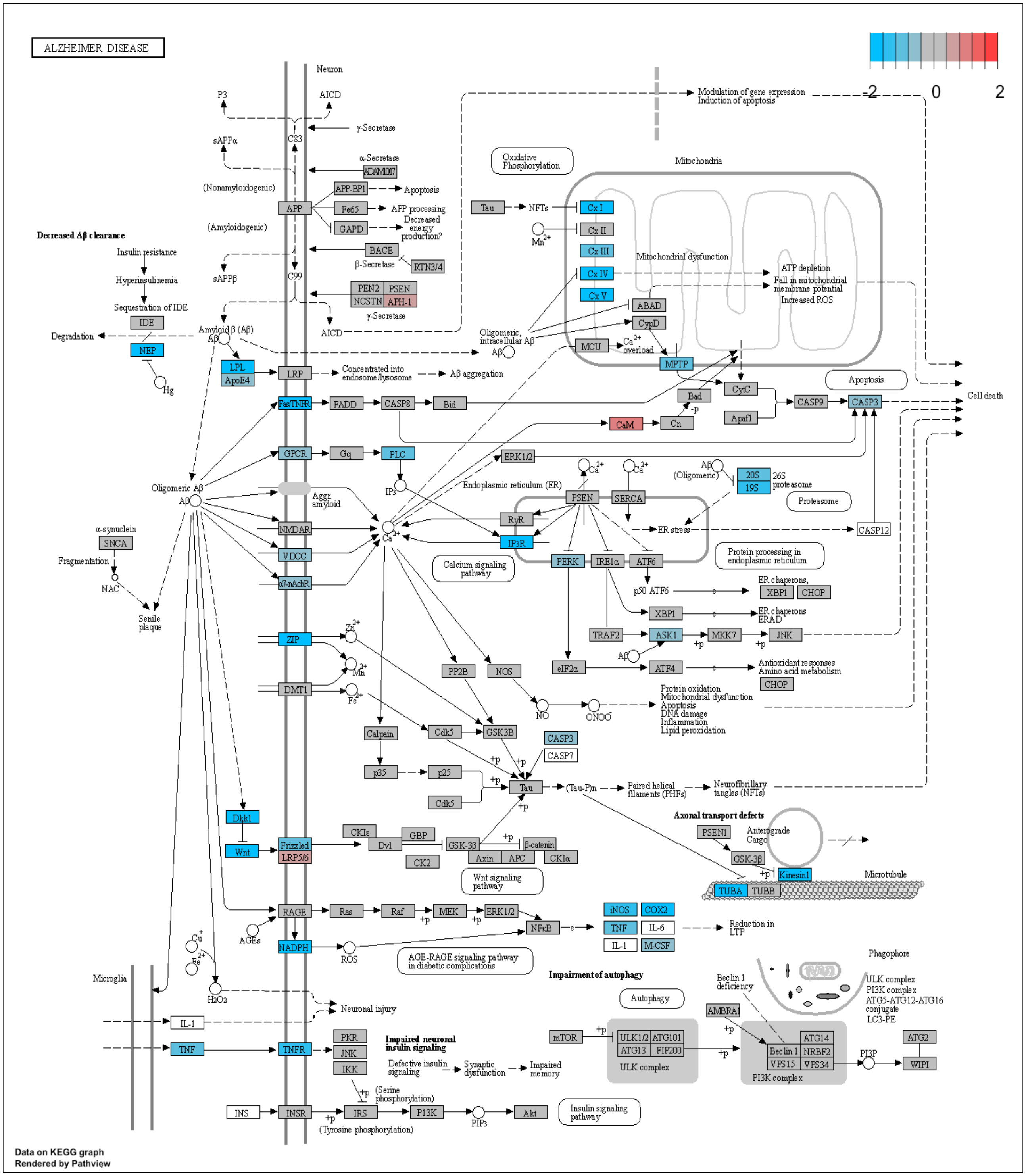


**Supplementary Figure 6. CBP KD Results in Widespread Downregulation of AD Pathway genes.** KEGG pathway view of AD-related differential gene expression pattern between and CBP KD and CTRL RNAi in APP$^{Dup}$ neurons generated in Pathview. Genes are color-coded based on expression from highest in KD (red) to lowest in KD (blue).

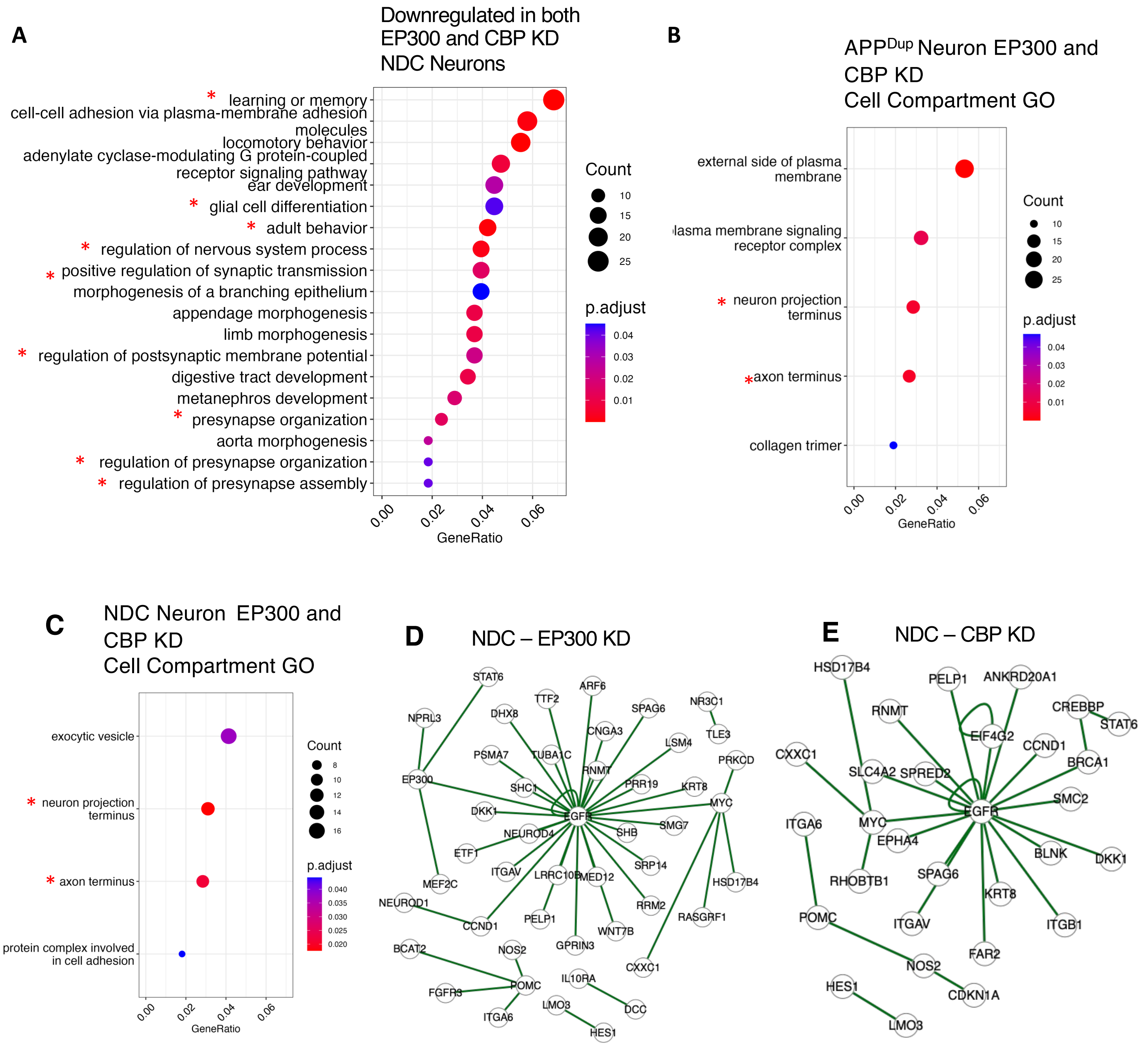


**Supplementary Figure 7. Cell compartment gene ontology analysis of EP300/CBP KD in APP^Dup and NDC neurons and esyN analysis of EP300/CBP KD in NDC neurons. A)** Bubble plot depicting gene ontology (GO) terms identified by ClusterProfiler as significantly enriched in the set of genes downregulated in both EP300 and CBP KD in NDC neurons**.** Neuron-related terms are asterisked. **B, C)** Bubble plots representing top Gene Ontology (GO) terms identified as enriched in genes downregulated in both EP300 and CBP KD in APP^Dup neurons **(B),** and in genes downregulated in both EP300 and CBP KD in NDC neurons **(C)** Neuron-related terms are asterisked**. D, E)** EGFR is a central interactor of genes significantly downregulated by P300 and CBP KD in NDC neurons. BioGRID-identified genetic interactions depicted by EasyNetworks (esyN) of genes significantly downregulated by EP300 **(D)** and CBP **(E)** KD.

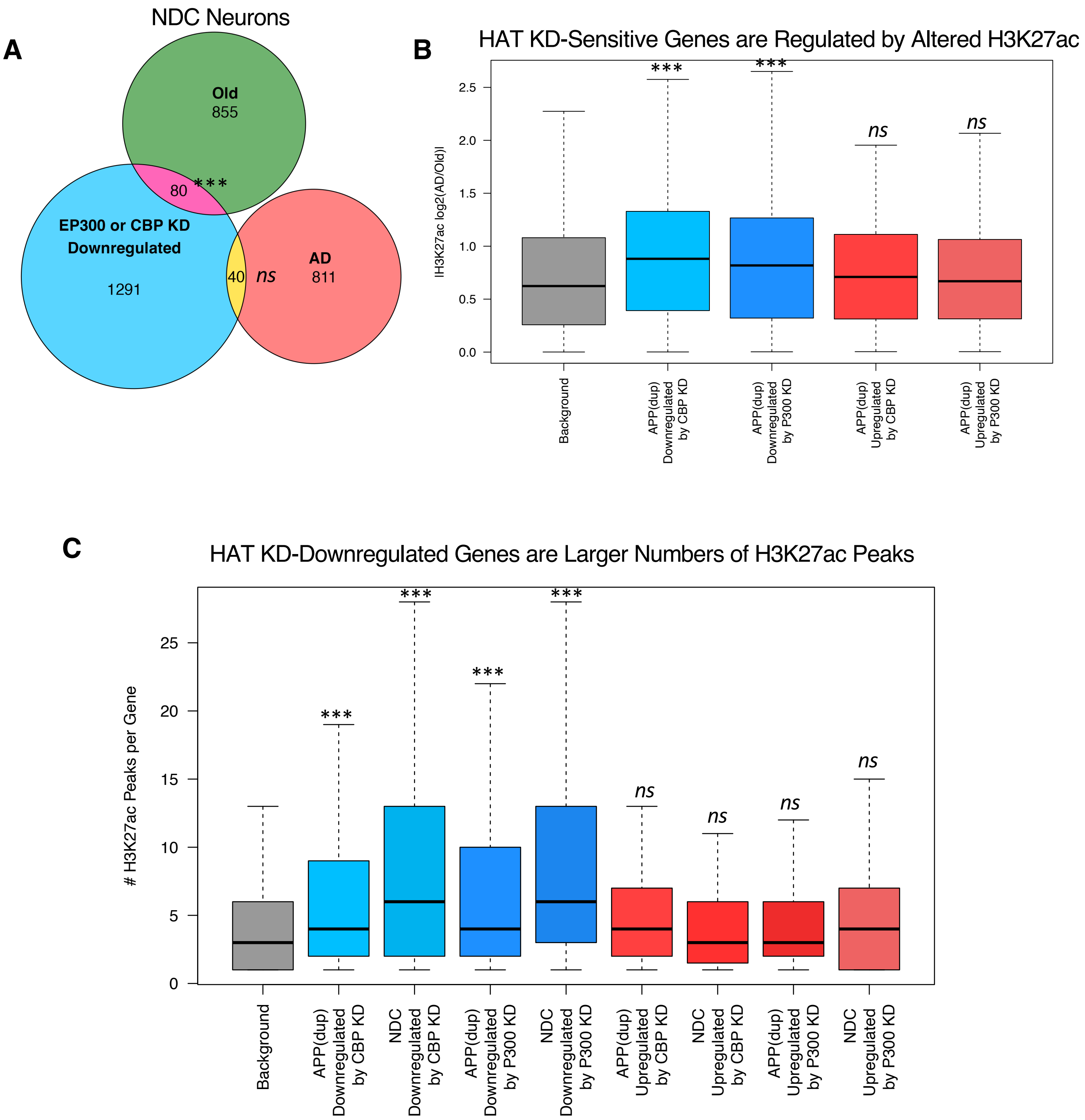


**Supplementary Figure 8. EP300 and CBP-regulated genes are subject to altered H3K27ac-defined enhancer regulation in AD and Old postmortem brain tissue.**
**A.** Significant overlap was observed between NDC EP300/CBP KD-Downregulated genes and Old-associated genes (Jaccard index = 80/2226, p =6.65e-06), but not between and NDC EP300/CBP KD-downregulated genes and AD-associated genes (Jaccard index = 40/2142, p = 0.75). Hypergeometric statistical testing was performed using the SuperExactTest R package. **B.** EP300- and CBP-regulated genes in NDC neurons are significantly associated with either AD-specific or Old-specific H3K27ac enrichment in human postmortem tissue, when contrasted with a control group of HAT-insensitive background genes that have nearby H3K27ac enrichment in the same brains. Statistical significance calculated using permutation test. **C.** Genes sensitive to HAT KD have more H3K27ac peaks / enhancers in human postmortem brain than expected by chance. Total number of peaks in both AD and Old samples per gene was counted and averaged for the sets of genes regulated by each enzyme in both $APP^{Dup}$ and NDC neurons. Statistical significance calculated using permutation test.

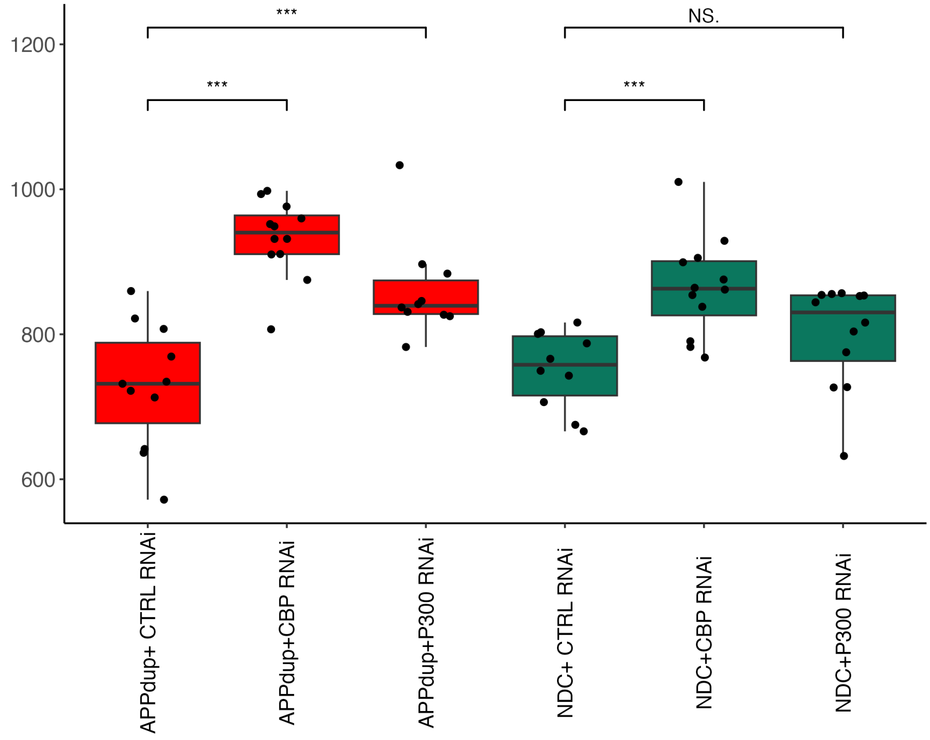


**Supplementary Figure 9. EP300 KD and CBP KD Effect on BACE1 expression in APP$^{Dup}$ and NDC neurons.** EP300/CBP KD results in markedly increased BACE1 expression in APP$^{Dup}$ neurons, but results in a smaller increase (CBP) or no significant change (EP300) upon KD in NDC neurons. Normalized read counts obtained through RNAseq.

Data Below only for safekeeping / organization – not mentioned

### Comparing Present Study to Nativio et al 2018

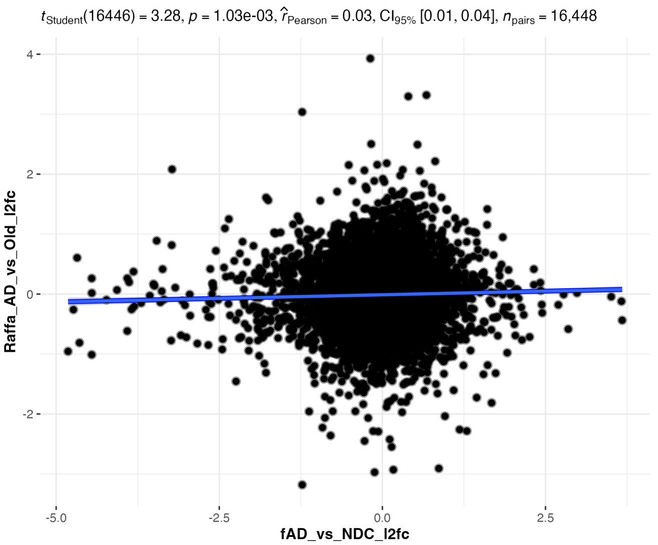


**A.** No correlation between (fAD vs NDC) vs. (Nativio AD vs Old)

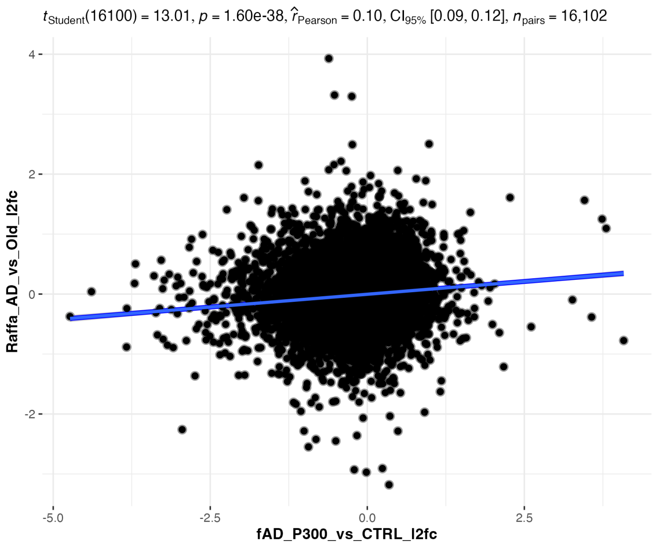


**B.** Weak correlation (R = 0.10) between (P300 vs CTRL KD) vs. (Nativio AD vs Old)

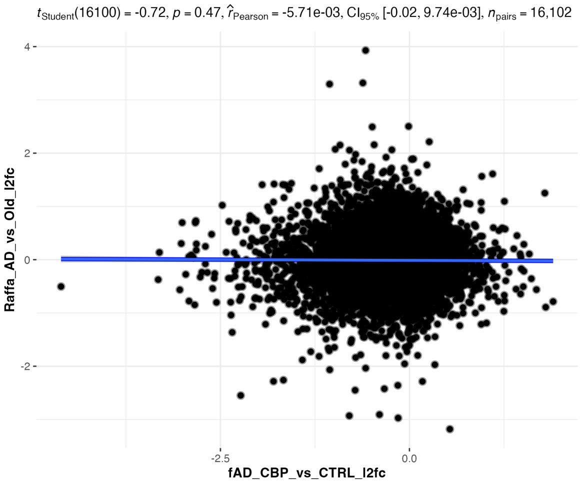


**C.** No correlation (R = 0.10) between (CBP vs CTRL KD) vs. (Nativio AD vs Old)

In our previous study, we reported that H3K27ac, an activating mark, is more abundant in the brains of AD patients relative to non-demented controls, and hypothesized that this reflected a dysregulated epigenomic state that might lead to expression of disease-promoting genetic pathways. In the context of heterogenous brain tissue, the balance of H3K27ac-mediated effect may indeed shift toward….

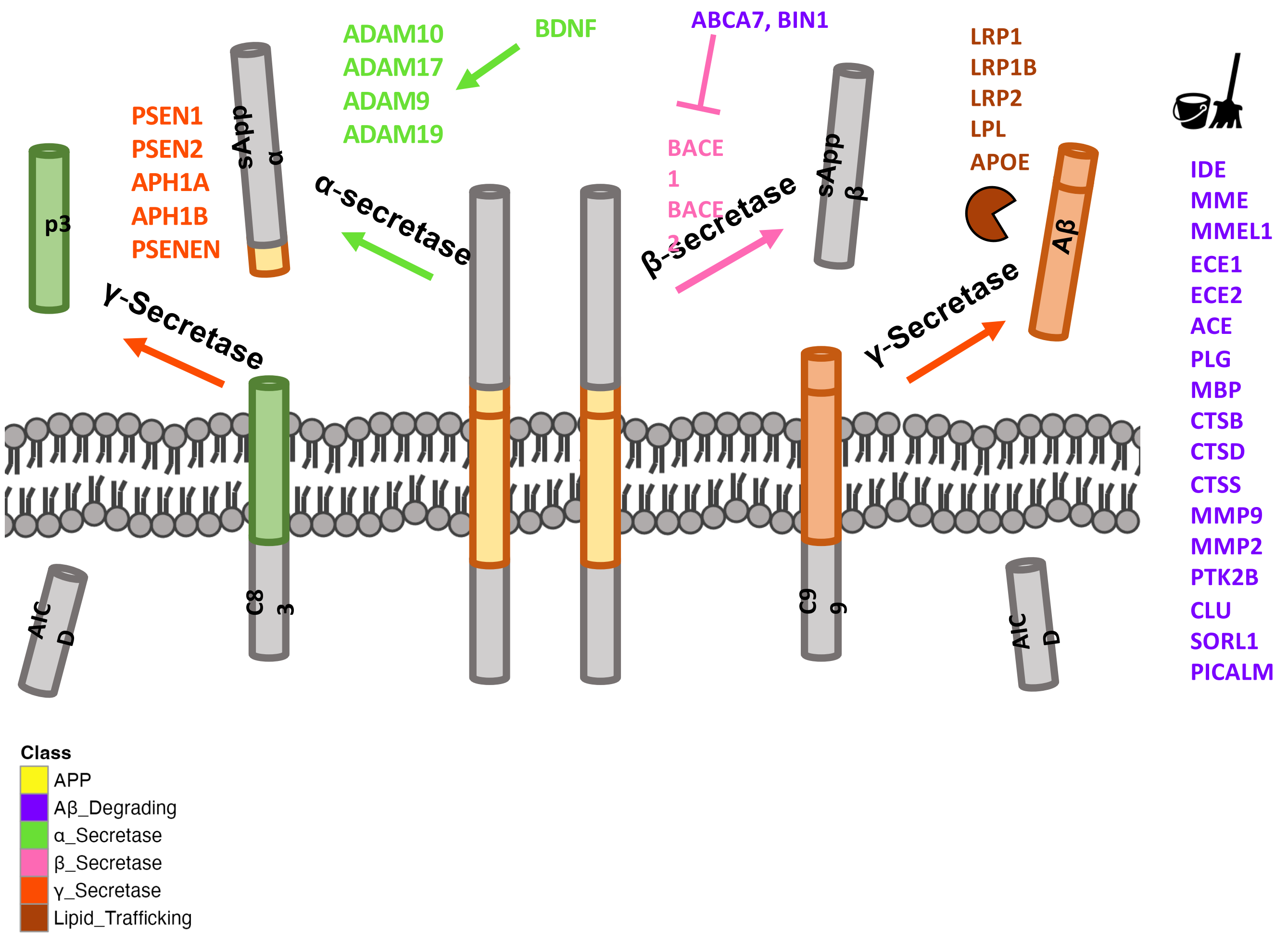

ADAM10
ADAM17
ADAM9
ADAM19
BDNF
ABCA7, BIN1
LRP1
LRP1B
LRP2
LPL
APOE
PSEN1
PSEN2
APH1A
APH1B
PSENEN
sApp α
p3
α-secretase
BACE 1
BACE 2
β-secretase
sApp β
Aβ
γ-Secretase
γ-Secretase
IDE
MME
MMEL1
ECE1
ECE2
ACE
PLG
MBP
CTSB
CTSD
CTSS
MMP9
MMP2
PTK2B
CLU
SORL1
PICALM
AIC D
C8 3
C9 9
AIC D
Class
APP
Aβ_Degrading
α_Secretase
β_Secretase
γ_Secretase
Lipid_Trafficking

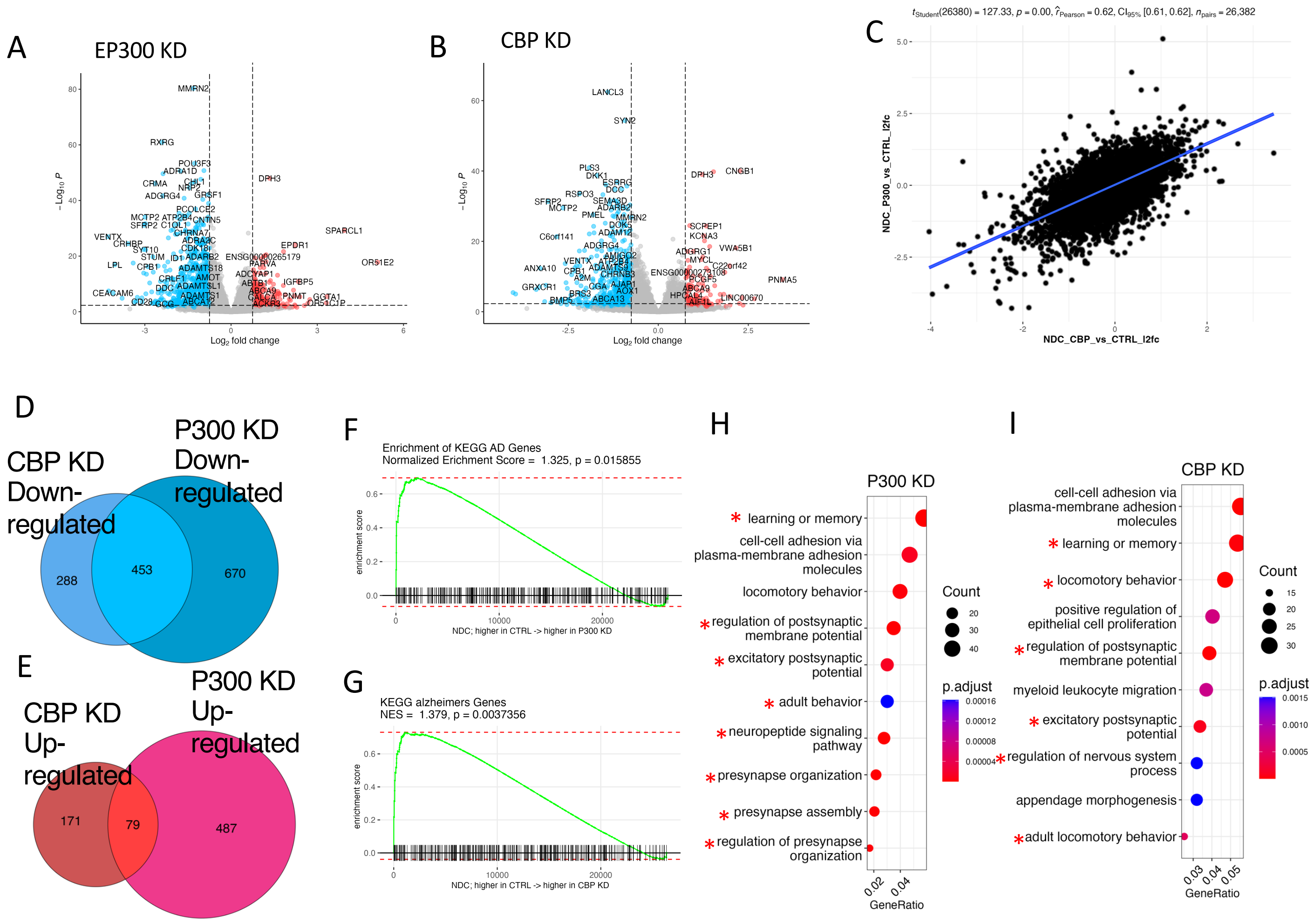

**Supplementary Figure 4. Transcriptional effects of EP300/CBP KD in NDC neurons. A, B)** EP300/CBP KD results in widespread transcriptional downregulation in NDC neurons. Volcano plot of gene expression changes between EP300 KD vs CTRL siRNA **(A)** and CBP KD vs CTRL siRNA **(B)** in NDC neurons. **C)** EP300 and CBP KD evoke generally similar transcriptional responses. Scatterplot depicting relationship of gene transcriptional changes between EP300 KD and CBP KD ($r_{pearson}$ = 0.62, *p* = 0.00) in NDC neurons. **D,E)** Venn diagram showing overlaps of gene expression changes between P300 and CBP KD in NDC neurons. Jaccard index of 453/1411, *p* < 0.05, hypergeometric test, calculated for overlaps between downregulated genes **(D).** Jaccard index of 79/737, *p* = 0.05, hypergeometric test, calculated for overlaps between upregulated genes **(E)**. **F, G)** KEGG-identified AD-associated genes are positively transcriptionally controlled by EP300 **(F)** and CBP **(G)** in NDC neurons. Gene set enrichment analysis (GSEA) plot depicting placement of KEGG AD genes in transcriptome ranked from highest expressed in CTRL RNAi to highest expressed in EP300 **(F)** and CBP **(E)** KD in NDC neurons. **H, I)** Bubble plot depicting gene ontology (GO) terms identified by ClusterProfiler as significantly enriched by EP300 **(H)** and CBP **(I)** KD in NDC neurons.

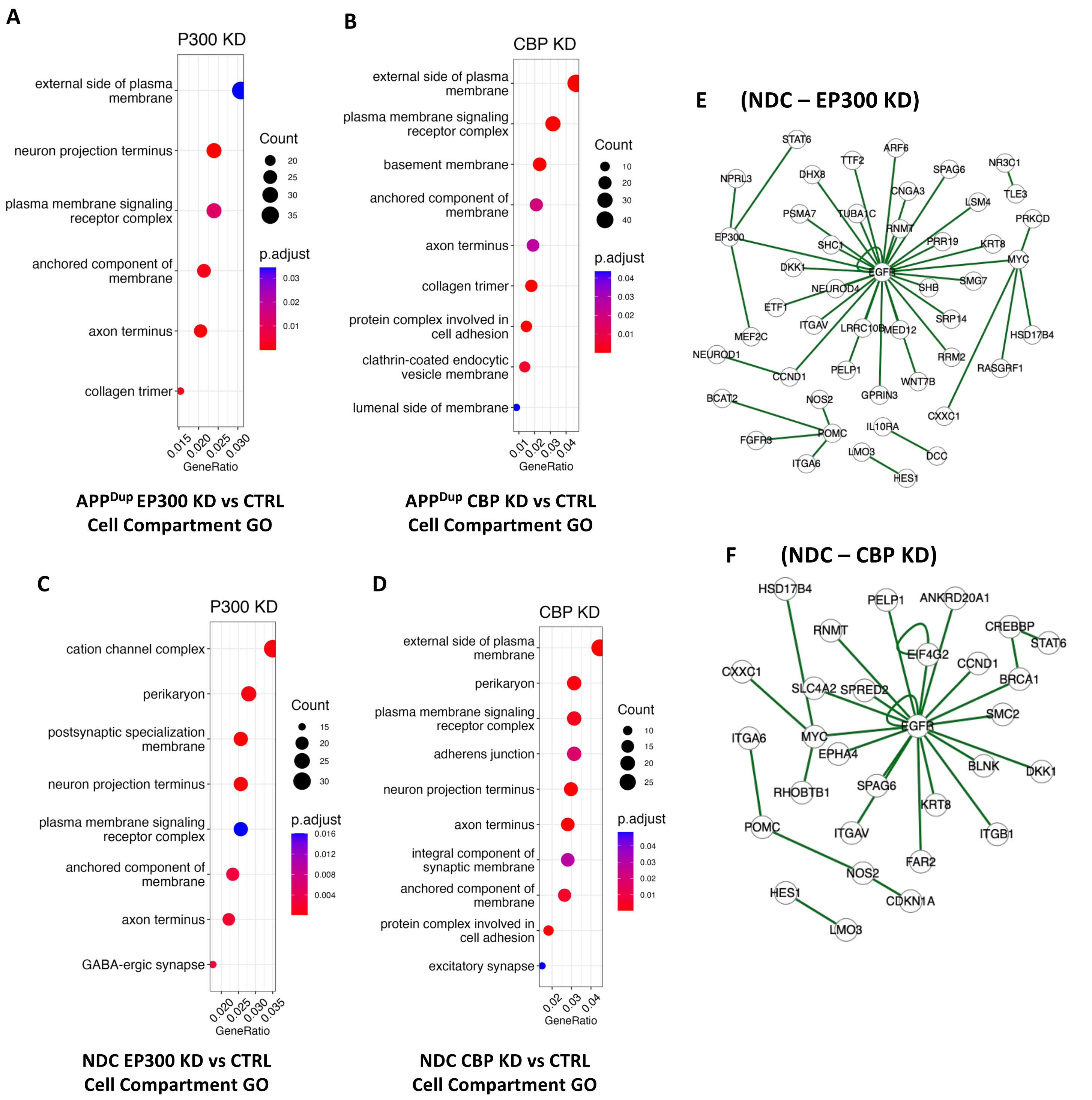


**Supplementary Figure 5. Cell compartment gene ontology analysis of EP300/CBP KD in APP$^{Dup}$ and NDC neurons and esyN analysis of EP300/CBP KD in NDC neurons. A-D)** Bubble plots representing top gene ontology (GO) terms identified as enriched in EP300 KD-downregulated genes in APP$^{Dup}$ neurons **(A)**, CBP KD-downregulated genes in APP$^{Dup}$ neurons **(B),** EP300 KD-downregulated genes in NDC neurons **(C)**, and CBP KD-downregulated genes in NDC neurons **(D)**. **E, F)** EGFR is a central interactor of genes significantly downregulated by P300 but not CBP KD in NDC neurons. BioGRID-identified genetic interactions depicted by EasyNetworks (esyN) of genes significantly downregulated by EP300 **(E)** and CBP **(F)** KD.
